# Sequence-Defined Digital Bottlebrush Polymers for Programmable Oligonucleotide Delivery

**DOI:** 10.64898/2026.05.09.723525

**Authors:** Jiachen Lin, Tingyu Sun, Yun Wei, Chenyang Xue, Guobin Xu, Peiru Chen, Yuyan Wang, Shaobo Yang, Chloe Cavazos, Caroline Shen, Allison Wang, Alex Wang, Ke Zhang

## Abstract

Oligonucleotide therapeutics hold transformative potential, yet their clinical translation is hindered by delivery barriers, including rapid renal/hepatic clearance and poor organ specificity. Bottlebrush polymers conjugates have emerged as a promising vector to address these limitations, but conventional architectures with uniform backbones can only achieve an unmodifiable, rigid biodistribution profile. Here, we report a library of sequence-defined “digital” bottlebrush polymers, precisely engineered with controlled placements of chemical motifs that modify physiochemical properties - including lipids, cholesterol, and cationic groups - along a polyphosphodiester backbone. Systematic evaluation of the digital bottlebrush polymer library reveals distinct structure-property relationships and enables organ-biased systemic delivery to several traditionally difficult-to-reach tissues, including muscle and skin. In a mouse model of rheumatoid arthritis, a single dose of a spleen-homing polymer-conjugated antisense oligonucleotide targeting TNF-α achieves potent knockdown and drives full functional recovery. These findings establish a versatile design framework for tailoring bottlebrush polymers to specific therapeutic applications.

## Introduction

Oligonucleotides have emerged as a powerful modality capable of modulating gene expression, splicing, post-transcriptional modifications, and non-nucleic acid target binding^1-4^, enabling therapeutics development for a broad spectrum of diseases. Despite their versatility, clinical success remains limited to a small number of concentrated disease settings in part due to poor delivery to extrahepatic organs and tissues, cell types, and subcellular organelles^5, 6^. Current delivery strategies, including chemical modifications^7^, antibody^8^ or ligand conjugation^9^, and nanoparticle systems such as lipid nanoparticles and polyplexes, are frequently constrained by toxicity, infusion reactions, immunogenicity, drug stability^10-12^, and limited delivery outside of a few targetable organs following systemic administration^13-18^. These limitations underscore the need for safe, non-immunogenic, and physiochemically stable delivery platforms that can be rationally programmed to satisfy disease-specific delivery requirements with regard to organ tropism and cell uptake^18^.

Towards this goal, we present a novel class of bottlebrush polymers with a sequence-defined “digital” backbone synthesized via solid-phase phosphodiester chemistry, which enables precise control over monomer identity, number, and spacing. Upon successful backbone synthesis, each monomer is derivatized with a polyethylene glycol (PEG) side chain to complete the bottlebrush architecture (Figure 1). A library of digital bottlebrushes containing systematically varied distributions of hydrophobic (C18, cholesterol) and cationic (spermine) monomers within the backbone was constructed. This modular structure offers a framework to subtly tune the physiochemical properties of the polymer. Even though the variations among the bottlebrush polymers may be small given their predominant PEG composition (>95% by mass), we hypothesize that the slight differences in how these materials interact with the plasma and cells can be “magnified” by a cumulative, “multi-pass” effect afforded by the extended plasma exposure of the bottlebrush polymer, driving distinct biodistribution patterns. In this regard, the digital bottlebrush polymers provide an alternative mechanism for engineering tissue tropism to target antibodies: where antibodies bind to known receptors for targeted delivery, the bottlebrush polymers achieve biased uptake in selected organs and tissues based on their physiochemical characteristics for passive delivery. The therapeutic oligonucleotide is covalently conjugated to the backbone of the bottlebrush polymer, which not only enhances the plasma pharmacokinetics (PK) of the oligonucleotide and extrahepatic accumulation, but also protects the oligonucleotide from unwanted nucleic acid-protein interactions, which can lead to side effects such as immune system activation and coagulopathy.

**Figure 1.**
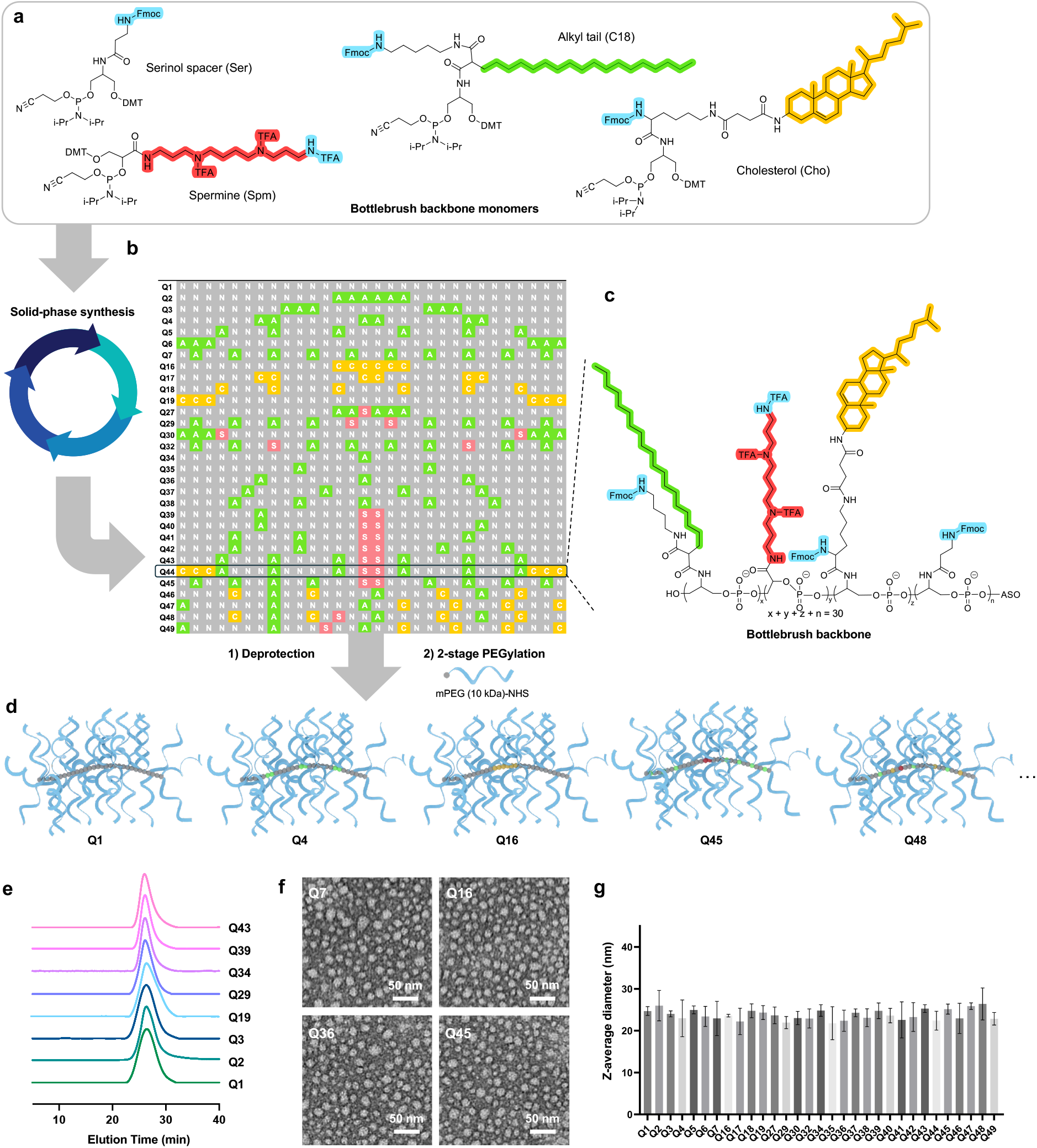
Synthesis and characterization of sequence-defined bottlebrush polymers. (A) Chemical structures of phosphoramidite monomers used for backbone synthesis. (B) Backbone library design. Each row in the table represents a unique bottlebrush backbone sequence, with sequence identifiers Q1-Q49. **N** (grey): serinol monomer; **A** (green): C18 alkyl monomer; **C** (yellow): cholesterol monomer; **S** (red): spermine monomer. (C) A generalized chemical structure of the sequence-defined bottlebrush polymer backbone. (D) Schematic representation of the final bottlebrush polymer architecture. (E) Aqueous size exclusion chromatograms of representative bottlebrush polymers, showing uniform molecular size (∼300 kDa M_n_) and polydispersity. (F) TEM images (negatively stained with 2% uranyl acetate of representative bottlebrush polymers. (G) Z-average hydrodynamic diameters of bottlebrush polymers determined by dynamic light scattering.

It was discovered that the bottlebrush polymer backbone significantly affects their cellular uptake rate^19-21^, with high-uptake samples exceeding lipofectamine 2k-assisted delivery despite being overall anionic. Importantly, by varying polymer backbone chemistry, substantial uptake in several conventionally hard-to-reach sites such as skeletal muscle, heart, and skin can be achieved, surpassing levels of state-of-the-art approaches such as targeting antibodies or Fab^22^. To demonstrate the utility of this platform, we identified a spleen-homing polymer (Q16) and applied it to the delivery of antisense oligonucleotides (ASOs) targeting tumor necrosis factor-alpha (TNF-α), a central inflammatory cytokine in autoimmune conditions such as rheumatoid arthritis (RA), Crohn’s disease (CD), and ulcerative colitis (UC)^23-25^. In a collagen-induced RA mouse model^26-28^, we demonstrated marked spleen ASO enrichment compared to free ASO (>2500-fold at 72 h post injection), efficient uptake by key immune cell populations in the spleen, and strong target engagement and functional recovery. Altogether, our results establish digital bottlebrush polymers as a programmable, generalizable platform with the potential to unlock oligonucleotide access to diverse organs and tissues once thought difficult or inaccessible.

## Results and Discussion

### Design and Synthesis of Digital Bottlebrush Polymer Library

The polyphosphodiester backbone of the bottlebrush polymer was assembled via a stepwise condensation process, incorporating four distinct phosphoramidite monomers (C18, Cho, Spm, and Ser), all of which were synthesized with a common serinol core (Figure 1A). The C18 (aliphatic chain) and Cho (cholesterol) phosphoramidites provide hydrophobicity to the final conjugate, while the Spm (spermine) phosphoramidite introduces two positive charges with each monomer. Both C18 and Cho are hydrophobic, but Cho adds biologically relevant membrane affinity and ordered packing, while C18 provides flexible hydrophobic stabilization. The unmodified serinol unit (Ser) acts as a spacer, allowing precise control over the spacing of other monomers and ensuring that the length of the final backbone polymer is independent of the number of modifiers present. A degree of polymerization of 30 is identified an optimal length based on our prior studies of polynorbornene-based bottlebrush polymers^19, 29^. All monomers contain an Fmoc- or trifluoroacetic acid (TFA)-protected primary amine, which, upon deprotection, is used to conjugate with a 10 kDa PEG side chain.

The monomers were synthesized by multi-step reactions ranging from 2 to 5 steps (Schemes S1-S4). Briefly, the C18 phosphoramidite was synthesized from octadecylsuccinic anhydride, which was grafted with mono-Fmoc-1,5-diaminopentane to incorporate a side-chain amine group. Subsequently, a mono-dimethoxytrityl serinol (DMT-serinol) was coupled by dicyclohexylcarbodiimide (DCC) and *N*-hydroxysuccinimide (NHS) chemistry. Finally, 2-cyanoethyl *N,N*-diisopropylchlorophosphoramidite was coupled to the remaining serinol hydroxyl group to finalize the C18 phosphoramidite (**5**). Similarly, starting with cholesterol amine, a two-step reaction was performed to yield cholesterol-NHS (**6**), which was reacted with Fmoc-Lys-OH to graft an amine group, producing cholesterol-Lys-OH (**7**). Cholesterol-Lys-OH was reacted with serinol-DMT and 2-cyanoethyl *N,N*-diisopropylchlorophosphoramidite to afford the Cho phosphoramidite (**9**). For the Spm phosphoramidite, spermine was mono-acylated with methyl (R)-2,2-dimethyl-1,3-dioxolane-4-carboxylate, followed by amine protection with trifluoroacetic anhydride to give spermine-prot-diol-TFA (**11**). After diol deprotection, 4,4-dimethoxytrityl chloride (DMT chloride) and 2-cyanoethyl *N,N*-diisopropylchlorophosphoramidite was reacted with the primary and secondary hydroxyl groups, respectively, producing the final Spm phosphoramidite (**13**). For the Ser (serinol) monomer, Fmoc-protected β-alanine was coupled with serinol using NHS/DCC, followed by DMT chloride substitution and phosphitylation to yield the final product (**16**). The chemical structures of all monomers and key intermediates are confirmed by ^1^H/^13^C NMR and mass spectrometry (Figures S9-29). The monomers were individually tested on a DNA synthesizer using standard coupling protocols, demonstrating sufficient coupling efficiencies (95%-98%).

Our primary objective in the digital backbone library design is to investigate how different monomer units, their numbers, arrangement, and combination, affect *in vitro* and *in vivo* characteristics of the final bottlebrush polymer. For the C18 monomers, we systematically increased the number of monomers from one to ten to evaluate the impact of hydrophobicity. Further, in the case of bottlebrushes with exactly six C18 monomers, we also examined the effect of spatial proximity from clustered to evenly distributed along the backbone (Figure 1B, Table S1). Similarly, we incorporated six units of Cho into the backbone and compared the effects of clustering vs. separation. To introduce positive charge, one or two Spm units were integrated into the C18 sequences, allowing us to assess the number and proximity effect of charge. Backbones with higher numbers of Spm monomers were successfully synthesized, but they were challenging to PEGylate and thus were excluded from the current library design. A library of 31 digital bottlebrush backbones were advanced to the next step, with overall yields between 20% to 50%. Following Fmoc removal with 20% piperidine in dimethylformamide (DMF), a two-step PEGylation process (first in aqueous buffer, then in DMF) using methoxy- and NHS-terminated 10 kDa PEG (mPEG_10k_-NHS) was carried out to derivatize backbone amine groups, achieving 94% average derivatization as determined by a 2,4,6-trinitrobenzene sulfonic acid assay (Figures 1B, C and D). The final bottlebrush polymers were purified by aqueous size-exclusion chromatography (SEC) to remove small molecule residues and excess PEG. Final purified polymers exhibit an M_n_ of approximately 300 kDa and uniform molecular weight distribution with polydispersity indices in the range of 1.06-1.19 for all samples (Table S2, Figure S2 and Figure 1E). Transmission electron microscopy (TEM) of samples stained with 2% uranyl acetate shows a spherical morphology with an average diameter of 20 ± 3 nm (Figure 1F and Figure S1). These measurements were corroborated by dynamic light scattering (DLS), which indicates approximately 23.7 ± 1.1 nm in hydrodynamic diameter (Z-average, Figure 1G). Collectively, these results show that the digital bottlebrush polymers are uniform in size and molecular weight, and thus their differences in biological interactions such as cell uptake rate and biodistribution should reflect the intrinsic physiochemical properties of the polymer.

### Backbone Structure of Digital Bottlebrush Polymers Strongly Dictates Cellular Uptake

To investigate the influence of the bottlebrush polymer backbone on their cellular uptake, three representative human cells, NCI-H358 (lung adenocarcinoma cell), HepG2 (hepatoblastoma cell), and GM03989 (primary muscle fibroblast) were treated with 0.5 µM of bottlebrush polymers for 4 h before confocal microscopy and flow cytometry analysis (Figures 2A, B and Figures S3A, B and C). Interestingly, the bottlebrush polymers yielded a large range in cell uptake rates, with the highest-to-lowest ratio being 50 to 300-fold across the cell lines tested. Remarkably, high-uptake particles like Q45 and Q29 readily exceed the uptake levels of Lipofectamine2k polyplexes in NCI-H358 cells while having an overall anionic charge. Further, the ranking of uptake is generally similar across the three cell types: high-uptake particles (e.g. Q45) remain high for all cells, while Q1 (Ser monomer only) consistently shows the lowest cell uptake, suggesting similar mechanism for cell uptake independent of cell type. Notable exceptions include Q41, Q2, and Q3: Q41 shows unusually high uptake in GM03989 while Q2/3 show high uptake in NCI-H358 cells, possibly due to cell-specific ligand-receptor interactions.

**Figure 2.**
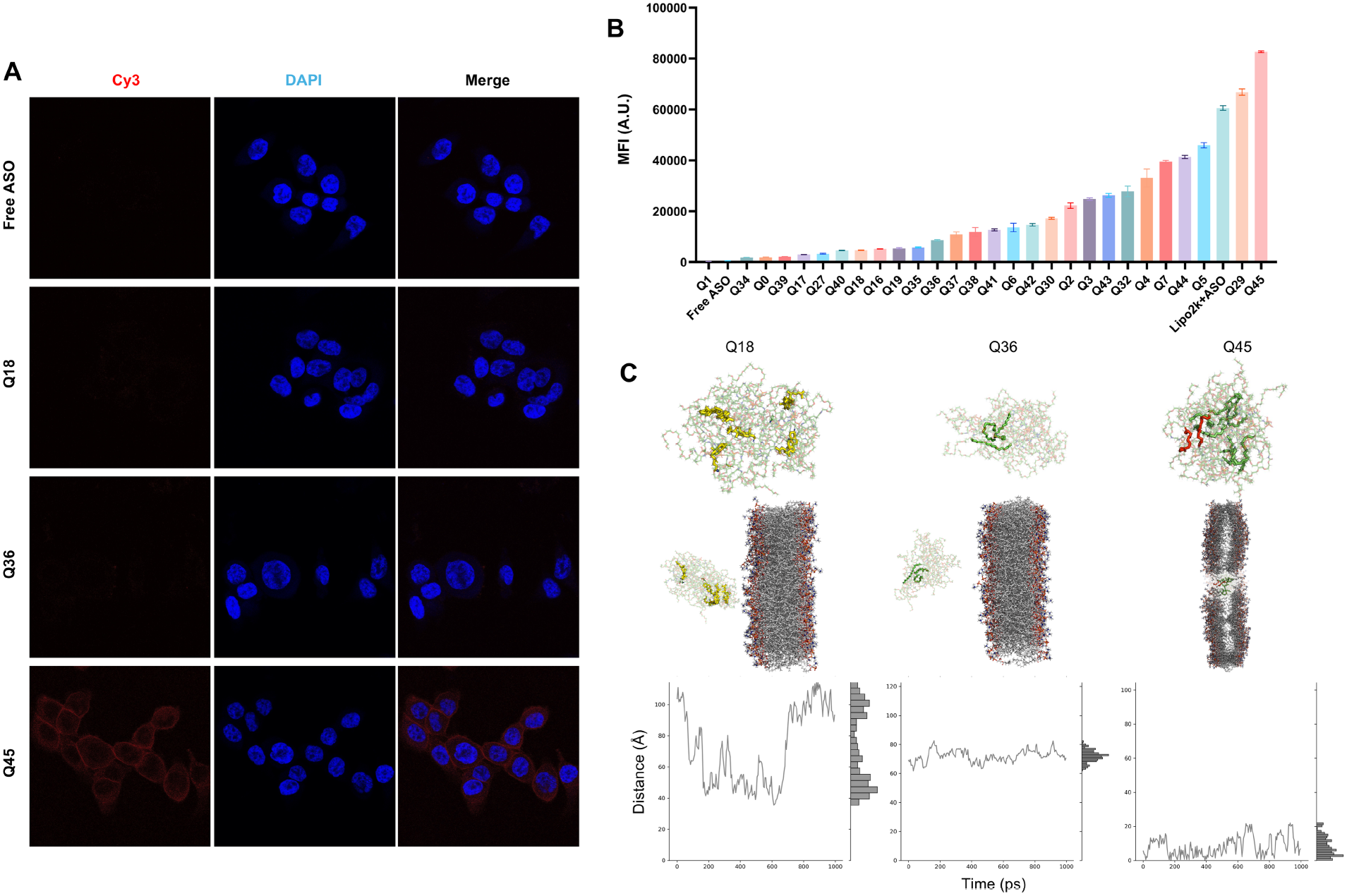
Cellular uptake and membrane interactions of bottlebrush polymers. (A) Confocal microscopy images of NCI-H358 cells treated with selected Cy5-labeled bottlebrush polymer samples (500 nM) for 8 h. (B) Mean florescence intensity (MFI) of NCI-H358 cells treated with bottlebrush polymers as determined flow cytometry. (C) Molecular dynamics simulations of three representative bottlebrush polymers in the presence of a model lipid bilayer showing membrane interaction dynamics.

Examining cellular uptake across the entire library, two main patterns can be gathered. First, cell uptake generally increases with the number of hydrophobic C18 monomers (seen in the series Q34-38, Q7). In addition, dispersed distribution of C18 monomers is more favorable vs. clustered distribution for high cell uptake (series Q2-6). These effects can be rationalized by the overall hydrophobicity of the polymer, which enhances the interaction with cells’ plasma membrane. Close placement of C18 with small spatial separation promotes self-interactions among the lipid tails, which reduces their collective hydrophobicity and increases overall solubility in water, leading to decreased cell membrane interactions. In fact, polymers with six clustered C18 (Q2/3) show less cell uptake than those with fewer C18 but with wider monomer separation (e.g. Q36-38). Notably, no straightforward trend was observed for the Cho series (Q16-19), which, like the C18 series, also contain six hydrophobic monomers per polymer. The second main pattern involves the Spm monomers, which can significantly enhance cell uptake, likely due to charge attraction with the anionic plasma membrane. In contrast to C18, clustering Spm monomers have a more pronounced effect in promoting cell uptake (series Q32, 29, 45) than separating them. This result indicates that concentrated positive charges have a larger effect than diffused charge of the same amount, and the clustered positive charges can exert a potent effect even when the molecule is net anionic. When the charge becomes too diffuse, the “sticky patch” effect is lost. With this hypothesis, it follows logically that the Q45 polymer, featuring eight evenly distributed C18 units and two adjacent Spm monomers, achieved the highest uptake across all cell lines. To further confirm these findings, Cy3-labeled Q18, Q36, and Q45 were selected to represent low-, medium-, and high-uptake polymers for confocal microscopy analysis using NCI-H358 cells treated with 0.5 µM polymer for 4 h. While Q18 showed minimum polymer-associated signals, Q36 and Q45 displayed apparent cell internal signals indicative of endocytosis. Further, for Q45, elevated signals were observed in the periphery of the cell, suggesting a high concentration of membrane-associated polymers.

To further investigate the initial polymer-cell membrane interaction leading to internalization, molecular dynamics (MD) simulations were conducted on the same representative polymers (Q18, Q36, and Q45) to probe the affinity of each polymer with a dipalmitoylphosphatidylcholine (DPPC) lipid bilayer (Figure 2C). The z-direction distance (perpendicular to the membrane surface between the center of mass of the polymer and the interface of the two membrane leaflets were recorded after two stages of 200 ns equilibration. Q45 was observed to maintain close proximity (0-2 nm) to the membrane surface, indicative of strong attractive interaction and partial bilayer embedding. Q36, with fewer C18 units and no Spm units, localized at a greater distance of 6-8 nm, suggesting weaker interactions, while Q18, composed six evenly spaced cholesterol units, exhibited widely fluctuating distances, reflecting short and weak membrane association. This affinity hierarchy (Q45 >Q36 >Q18) is closely aligned with the flow cytometry and confocal microscopy data. Collectively, these results highlight the crucial role of monomer distribution, along with their number and type, in governing membrane interaction and cellular internalization.

### Plasma PK, Biodistribution, and Innate Immune System Activation

Next, the impact of bottlebrush polymer backbone on the plasma PK and biodistribution was investigated using C57BL/6 mice and Cy5-labeled bottlebrush polymers. Mice received 10 nmol of the polymer via tail vein intravenous (i.v.) injection and blood samples were collected over a 72-h period to assess plasma PK. At the study endpoint, major organs including brain, heart, liver, spleen, lung, kidney, muscle, and skin were harvested for quantitative biodistribution analysis. In addition to analyzing trends across the various bottlebrush backbone designs, plasma PK/biodistribution of the polymer is also compared to that of a free phosphodiester DNA ASO^30^.

All bottlebrush polymers demonstrated substantially prolonged circulation with 4.1%-27.3% of injected dose retaining in plasma after 24 h, attributed to reduced renal clearance due to the large molecular size (Figure 3A). In contrast, free ASO showed less than 0.1% in circulation after 24 h. Analysis of plasma PK reveals several structure-exposure relationships for the digital bottlebrush library. Increasing Cho content tend to reduce systemic exposure (area under the curve, AUC_∞_), likely due to enhanced hepatic or reticuloendothelial clearance. However, if Cho monomers are not grouped together, high systemic exposure can be retained. In contrast, higher proportions of Ser monomers were associated with prolonged half-life (t_½_), indicating that anionic backbone minimize nonspecific binding and confer greater circulatory stability. Interestingly, the Spm monomer produced only a modest effect in overall half-life (small increase), even though it has a large positive effect in cell uptake, indicating a limited influence on circulation despite its cationic nature (Supplementary Table 3).

**Figure 3.**
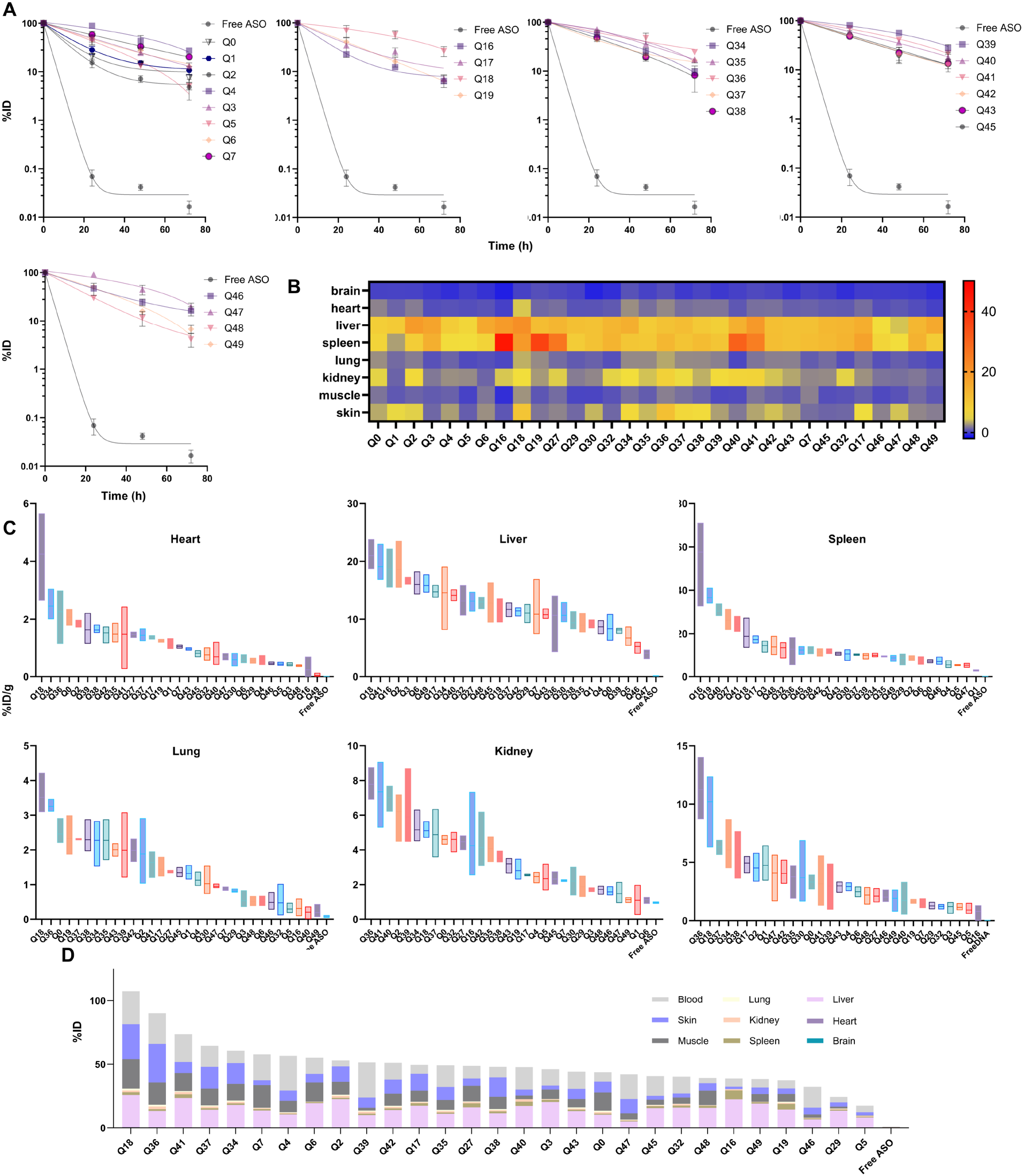
Quantitative plasma pharmacokinetics and biodistribution. (A) Plasma pharmacokinetics measured over 72 h following intravenous administration. (B) Heatmap representation of % injected dose per gram of tissue (%ID/g) across major organs for all tested bottlebrush polymers 72 h after injection. (C) Biodistribution bottlebrush polymers ranked by organ from high to low. (D) Estimated total retained polymers calculated from organ weights and %ID/g for each organ, shown as a stacked bar chart.

Next, we evaluated the biodistribution of the bottlebrush polymers, first qualitatively by *in vivo* imaging (IVIS) of excised major organs and then quantitatively by fluorescence measurements using homogenized tissues. It is evident that free oligonucleotide is rapidly cleared from the animal, with nearly no detected signal outside the kidney (Figure S4). In contrast, all bottlebrush polymer samples exhibited significantly increased signals across major organs (except the brain). A heatmap derived from the quantified organ-level biodistribution data revealed distinct accumulation patterns across tissues (Figure 3B). For each organ, polymer samples were ranked from high to low uptake to reveal trends (Figure 3C). Q16 demonstrated the highest accumulation in the spleen, with 57% injected dose per gram of tissue (%ID/g), which is over 10-fold that of the least spleen-tropic polymer and >1000-fold greater than the free ASO. This selective spleen retention is attributed to the presence of Cho monomers along its backbone, as the spleen is highly responsive to changes in lipid content. This observation is consistent with prior studies showing Cho-rich nanoparticles preferentially interact with the mononuclear phagocyte system, particularly with splenic macrophages^31^. An interesting discovery is that the clustering of Cho monomers generates a much stronger (3-fold) spleen-tropic effect than isolated Cho (series Q16-19) when identical numbers of Cho monomers are present.

This “backbonomics” approach can be a powerful technique for identifying nanoparticles capable of passive enrichment in traditionally difficult-to-reach tissues. For example, Q18 exhibited preferential accumulation in muscle tissues, with average muscle uptake at 2.6 %ID/g. Given that muscle accounts for approximately 35-40% of the animal’s wet mass, nearly a quarter (23%) of total injected dose accumulate in muscle tissues with Q18 after i.v. administration. This level of muscle tropism is markedly higher compared to other muscle-targeted approaches such as transferrin receptor-binding antibodies/Fab and integrin receptor-targeting peptides^32^. While the precise mechanism targeting remains to be investigated, polymers with high muscle uptake appear to share several features, including intermediate hydrophobicity, absence of cationic components, and long plasma retention. Interestingly, the combinatorial backbone design allows more stringent delivery requirements than single-organ enrichment to be satisfied. For example, it is possible to identify backbone sequences that achieve selective uptake by skeletal muscle or cardiac muscle.

In this regard, Q5 exhibits ∼10-fold cardiac muscle selectivity while Q49 shows ∼20-fold skeletal muscle selectivity. Another example of unmet drug delivery need involves the skin, as efficient skin-biased biomarkers that enable active targeting have not progressed beyond proof of concept^33^. Here, Q36 is identified as the best skin-targeted sequence, with approximately a third (30.4%) of injected dose accumulating in the skin (12% ID/g). The high skin enrichment may be attributed to long blood circulation times enabling improved cell exposure in the small, terminal vasculature associated with the skin (passive targeting). Long blood circulation, in turn, is positively correlated with low to moderate cell uptake, low to intermediate hydrophobicity, and lack of positive charge. In addition to identifying polymers with the highest uptake in specific organs, it may also be of value assessing the ability to de-target potential dose-limiting organs while maintaining reasonable uptake in the targeted organs. For example, Q47 exhibits very low liver (4.1% ID/g) and spleen uptake (5.2% ID/g), demonstrating limited clearance by the mononuclear phagocyte system (MPS). However, its uptake in skin is approximately 50% of Q36, which is still impressive. With the current test library, uptake in the brain remains minimal across all samples, suggesting inability to cross the blood-brain barrier (BBB). However, limited exposure in the central nervous system (CNS) may be beneficial in minimizing CNS-related toxicity. Together, these results underscore the critical role of backbone arrangement in determining bottlebrush polymers’ *in vivo* fate and demonstrate that subtle changes in polymer sequence can yield distinct organ-specific targeting patterns (Figures 3B, C).

Plotting the per-organ biodistribution of each bottlebrush polymer sample in an aggregated bar chart, it can be seen that all bottlebrush polymer structures exhibited substantially greater retention in major organs compared to free ASO, which is almost entirely cleared (>99.8%) by 72 h. This reaffirms that the PEGylated bottlebrush architectures effectively reduce renal clearance and prolong systemic exposure (Figure 3D). In most bottlebrush polymers, the uptake in four organs (muscle, skin, blood, and liver) account for the majority (>90%) of total organ-associated materials. Notably, there is a large range in the clearance rate among the different bottlebrush polymers, from nearly no clearance (Q18) to 83% clearance (Q5), assuming that samples not detected in major organs are cleared. There do not appear to be easily discernible structure-property relationships governing tissue retention; possible molecular features that promote long tissue retention include separation of Cho monomers, lack of charged units (Spm), and having three to four separated lipid tails (C18).

To evaluate the immunocompatibility of the bottlebrush polymer library, blood samples were collected from C57BL/6 mice 24 h after i.v. injection and were analyzed for inflammatory cytokine signals (Figure S5C). Lipopolysaccharide (LPS) and PBS were used as positive and negative controls, respectively (0-100% activation). Across all bottlebrush variants, no significant upregulation of pro-inflammatory cytokines, including TNF-α, IL-6, and IL-1β were detected, suggesting that the polymers are free from unwanted innate immune activation and supporting its use as a therapeutic delivery platform.

### Spleen-Specific Delivery Using Q16 for TNF-α Silencing in a Rheumatoid Arthritis Model

To demonstrate the ability of digital bottlebrush polymers to serve as an organ-biased delivery platform, we selected Q16, which shows rapid and high uptake in the spleen, as a proof-of-concept using collagen-induced arthritis (CIA) mouse model, a widely adopted RA murine model. RA is a systemic autoimmune condition driven by TNF-α, which sits at the top of an inflammatory cascade, regulating the production of other cytokines (IL-1, IL-6, GM-CSF), chemokines, and adhesion molecules. The spleen is a key organ for monocyte/macrophage and dendritic cell (DC) activation, both of which are central producers of TNF-α. These immune cells not only excrete TNF-α into the bloodstream but can also traffic to the synovium to enhance local joint inflammation. Therefore, targeting TNF-α production in the spleen reduces the circulating inflammatory pool that ultimately affects the joints. This strategy was adopted in the clinical evaluation of IONIS-104838, a 20-mer gapmer ASO, in TNF-α inhibitor-naïve RA patients. The ASO showed target engagement preclinically but ultimately failed to meet clinical efficacy endpoints in Phase II clinical studies, which was postulated to stem from inadequate delivery to spleen-resident immune cells^2, 5, 6, 34^. While TNF-α monoclonal antibodies have been wildly successful in RA, they are limited by their extracellular mechanism of action, high dosing requirements in patients with elevated TNF-α load, and the possibility of developing of anti-drug antibodies (ADAs) in up to 30% of patients^35^. Therefore, oligonucleotides, with an orthogonal mechanism (suppressing TNF-α at the transcript level) can still be an important therapeutic option in RA, and a delivery strategy that enables more effective engagement of the cellular reservoirs of TNF-α could offer renewed potential for an oligonucleotide-based therapy.

We first evaluated the performance of a mouse TNF-α-directed gapmer ASO formulated as a Q16 conjugate *in vitro*. RAW264.7 cells were pre-treated for 4 h with Q16-ASO and then maintained in ASO-free media for 24 h before LPS stimulation. Despite the absence of continued drug exposure, TNF-α secretion measured 2.5 h post-stimulation was reduced by 58% compared with control (Figure 4D). These results support the concept that, unlike target neutralization with biologics, intracellular knockdown provides functional suppression that does not depend on maintaining constant high drug concentration in the vicinity of the cells.

**Figure 4.**
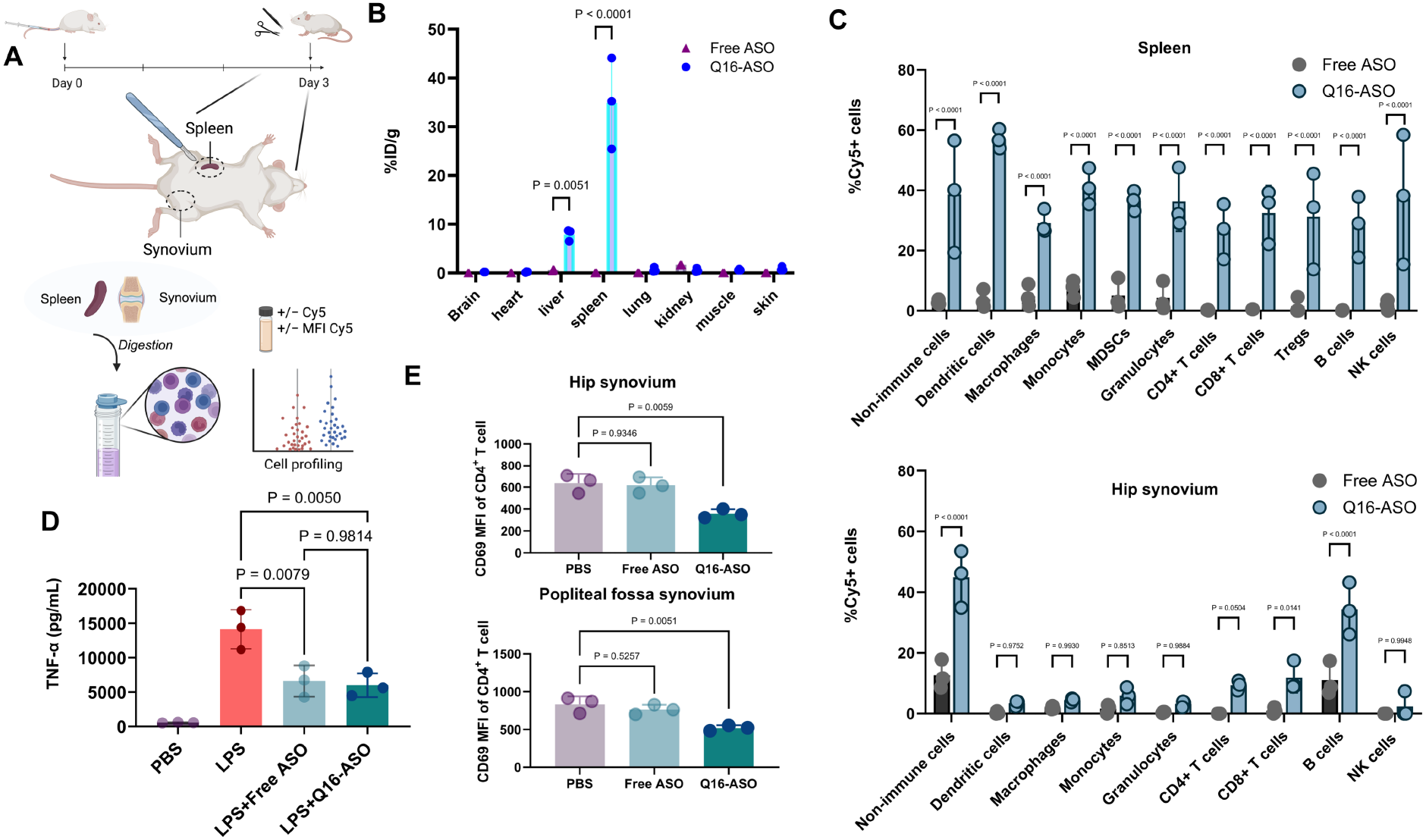
Q16-assisted spleen delivery of TNF-α-silencing ASO. (A) Treatment schedule and schematic. (B) Biodistribution of Q16-ASO in DBA/1J mice 72 h following i.v. injection. (C) Cy5^+^ immune and non-immune cell populations in the spleen and hip synovium 72 h following i.v. administration. (D) *In vitro* TNF-α knockdown in RAW264.7 cells. (E) CD69^+^ CD4^+^ T cells in the synovium 72 h following injection, demonstrating reduced CD4^+^ T cell activation. Statistical analysis was performed using one-way ANOVA.

To evaluate the ability of Q16 to deliver the TNF-α gapmer ASO to the spleen, Q16-ASO was administered i.v. to DBA/1J mice immunized with type II collagen in Complete Freund’s Adjuvant (CFA). Major organs were collected 72 h post-injection for biodistribution analysis, and primary splenocytes were isolated in parallel for flow cytometric profiling of immune cell subsets (Figure 4A). Biodistribution analysis showed that Q16-ASO retained strong spleen accumulation seen with the naked polymer, and ∼2,500-fold higher than free ASO (Figure 4B). Moreover, the organ distribution patterns of Q16-ASO closely matched those of the unconjugated polymer, indicating that the conjugate’s biodistribution is strongly dictated by the polymer scaffold. Flow cytometry further demonstrated markedly higher fractions of Cy5^+^ (ASO^+^) cells across major splenic immune subsets in Q16-ASO-treated mice, ranging from 4-to 100-fold increases compared to free ASO (Figure S4C), while the relative proportions of these cell populations remained unchanged (Figure 6A), suggesting that the conjugate does not induce gross immune perturbation such as cell depletion or expansion. Among splenic subsets, NK cells showed the highest uptake (mean fluorescence intensity, Figure S6B). In addition to the spleen, immune cells in the hip synovium were also profiled (Figure 4F). Consistent with the prolonged circulation of the polymer conjugate, the fraction of Cy5^+^ synovial immune cells were uniformly higher in the Q16-ASO group than in the free-ASO group, though to a lesser extent than in the spleen, underscoring the spleen-tropic nature of Q16. Notably, synovial CD4^+^ T cell activation was significantly reduced, evidenced by diminished CD69 expression (Figure 4E), suggesting potential pharmacological activity of Q16-ASO within the synovial compartment.

To evaluate the functional recovery associated with Q16-ASO treatment, DBA/1J mice were immunized with type II collagen twice (on Day 0 and then on Day 14). A single dose of either free ASO or Q16-ASO conjugate (13 mg/kg, approximately 20 nmol/animal, ASO basis) were administered i.v. on Day 7. Clinical signs of arthritis, particularly in the limbs, were monitored to assess disease severity, and animal body weight was tracked throughout the study (45 days) as an indirect indicator of systemic disease burden (Figures 5B, C). Treatment with Q16-ASO conjugates resulted in significantly reduced arthritis severity compared with both untreated control and free ASO, as indicated by lower arthritis scores (double blinded assessment). Further, while untreated and free ASO-treated groups exhibited progressive body weight loss, a common consequence of systemic inflammation^26, 28^, Q16-ASO-treated mice maintained constant body weight close to baseline levels, suggesting an overall reduction in disease stress and inflammation.

**Figure 5.**
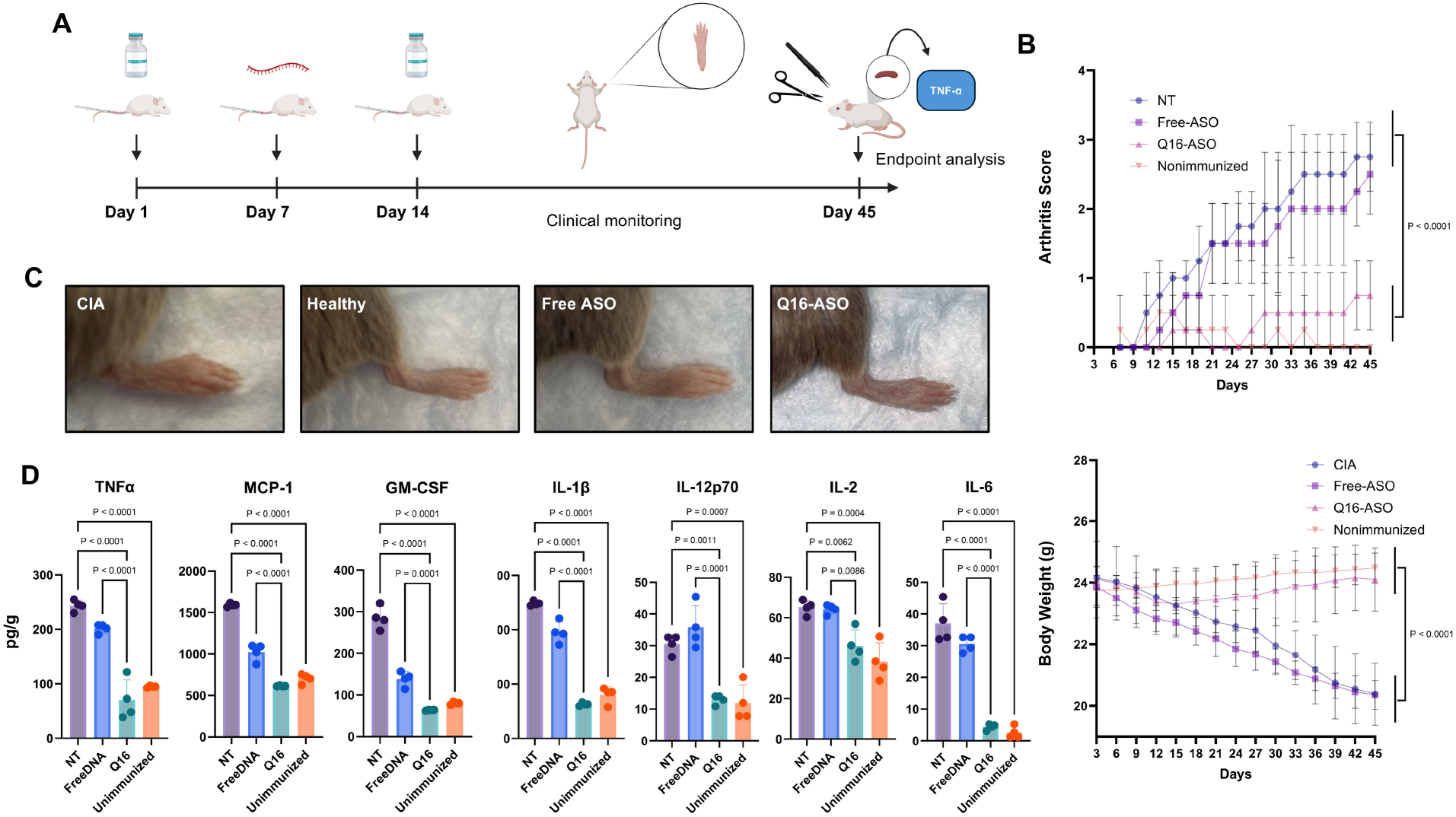
Therapeutic evaluation of Q16-ASO in a RA mouse model. (A) Schematic of disease phenotype induction and treatment schedule. (B) Arthritis score and animal body weight changes throughout the study. (C) Photographs of mouse hind paws showing visibly enlarged and inflamed paws for the CIA and free ASO treated groups, while Q16-ASO treated mice and healthy animals show normal paws with no visible edema. (D) Cytokine expression levels in the spleen tissues at endpoint (Day 45). Statistical analysis was performed using one-way ANOVA.

At the study endpoint (Day 45), mice were sacrificed for molecular and histological analyses. Homogenized spleen samples were evaluated for pro-inflammatory cytokines, including TNF-α, IL-6, IL-2, and IL-1β, while cryosectioned spleen tissues were processed for hematoxylin and eosin (H&E) staining. Because Q16 is designed to deliver TNF-α ASO directly to spleen-resident immune cells, we prioritized spleen cytokine quantification as the most proximal pharmacodynamic readout of target engagement. Cytokine analysis revealed significantly reduced levels of TNF-α and associated mediators in the Q16-ASO treatment group compared with both untreated and free ASO-treated groups (Figure 5D), close to healthy (unimmunized) levels. Histological evaluation further demonstrated that Q16-ASO-treated mice retained normal splenic architecture and cellular organization, in contrast to the disrupted and hyperplastic spleens observed in control mice (Figure S8). These findings indicate that spleen-targeted delivery of TNF-α ASO via Q16 enhances the pharmacological potency of the therapeutic oligonucleotide and mitigates RA progression effectively.

## Conclusion

In this study, we report the development of a sequence-defined digital bottlebrush polymer platform for organ-biased antisense oligonucleotide delivery. By systematically varying the type, number, and spatial arrangement of hydrophobic and cationic monomers, we established several structure-function relationships governing cellular internalization, plasma PK, and biodistribution. Notably, we identified Q16 as a spleen-selective polymer, which demonstrates high splenic accumulation and efficient immune cell delivery without inducing systemic immune activation. When conjugated with a TNF-α-targeting ASO, Q16 achieved potent therapeutic outcomes in a collagen-induced arthritis model, significantly reducing disease progression and pro-inflammatory cytokine expression. Together, our findings highlight the potential of digitally encoded bottlebrush polymers as a programmable and biocompatible platform for precision oligonucleotide delivery, paving the way for tailored therapies in extrahepatic contexts.

## Data Availability

All data are available in the main text or Supplementary Information. Source data are provided with this paper.

## Associated Content

Materials, experimental methods, synthesis and characterization of bottlebrush polymers, additional experimental data including sequences, RP-HPLC chromatograms and live mice imaging.

## Author Contributions

K.Z., J.L and T.S. devised the experiments and wrote the manuscript. J.L. conducted materials synthesis and material/biological characterization. T.S. conducted materials purification and material/biological characterization. Y.W. carried out biological analyses. All other authors contributed to material synthesis, biological assays, and MD simulation. All authors edited the manuscript.

This publication was made possible by the National Science Foundation (DMR award number 2503350), the National Institute of General Medical Sciences (1R01GM121612), the National Cancer Institute (4R42CA275425 and 5R01CA251730), and the Department of Defense Congressionally Directed Medical Research Programs (CDMRP MD230034).

## Acknowledgments

The authors thank Dr. Guoxin Rong from the Institute for Chemical Imaging of Living Systems at Northeastern University for assistance with flow cytometry, Dr. Shirin Kaboli from Boston Electron Microscopy Center (BE9MC). Dr. Jason Guo 500Hz and 700Hz NMR from NMR core facility.

## Competing Interests

The authors declare no competing financial interests.

## TOC

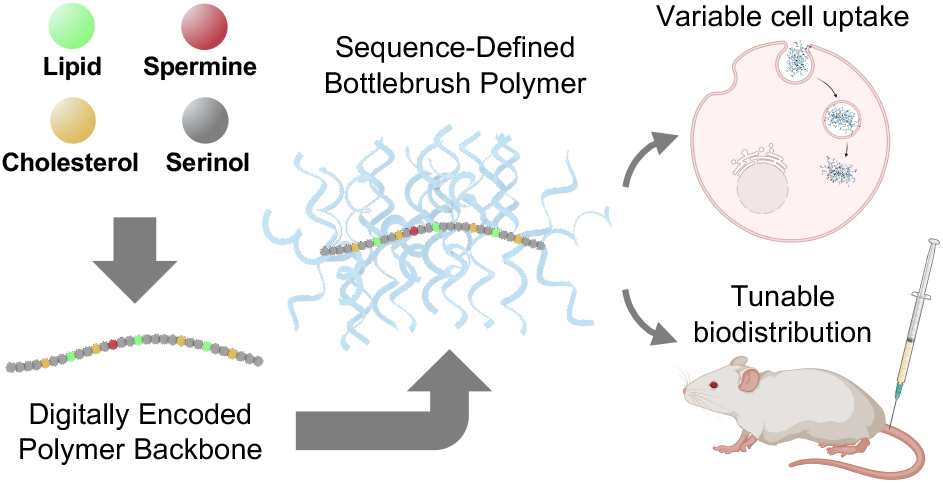

## Notes

### Competing Interest Statement

The authors have declared no competing interest.

