## Supplementary Information for "Sequence-Defined Digital Bottlebrush Polymers for Programmable Oligonucleotide Delivery"

#### Supplementary Methods

##### Material

Phosphoramidites and supplies for DNA synthesis were purchased from Glen Research Co. (Sterling, VA, USA). Methoxy polyethylene glycol (PEG) glutaramide succinimidyl ester ($M_n$=10 kDa) was purchased from Creative PEGWorks (Chapel hill, NC, USA). Human NCI-H358 lung cancer cells, human HEP-G2 cells, and GM03989 human DM1 fibroblast cells were purchased from American Type Culture Collection (Rockville, MD, USA). All chemicals were purchased from Fisher Scientific Inc. (USA), Sigma-Aldrich Co. (St. Louis, MO, USA), or VWR International LLC. (Radnor, PA, USA), and used as received unless otherwise indicated.

##### Reverse-Phase High-Performance Liquid Chromatography (RP-HPLC) and Size-Exclusion Chromatography (SEC)

Reversed-phase high-performance liquid chromatography (RP-HPLC) was performed on a Waters (Waters Co., Milford, MA, USA) Aqueous SEC measurements were performed on a Waters Breeze 2 SEC system equipped with Ultrahydrogel™ 1000, Ultrahydrogel™ 500 and Ultrahydrogel™ 250, 7.8Å×300 mm column and a 2998 PDA detector (Waters Co., Milford, MA, USA). Phosphate-buffered saline (PBS, pH 7.4) was used as the eluent running at a flow rate of 0.8 mL/min. DMF SEC analysis was performed on EcoSEC HLC-8320 SEC system (Tosoh Bioscience LLC, Tokyo, Japan) equipped with a TSKGel α-M 7.8×300 mm, 13 μm column and RI/UV-Vis detectors. The mobile phase is 0.05 M lithium bromide in HPLC-grade DMF, and samples were analyzed at a flow rate of 0.4 mL/min. SEC calibration was based on a ReadyCal kit of polyethylene glycol standards (PSS-Polymer Standard Service-USA Inc., Amherst, MA, USA). The kit covers an $M_n$ range from 232 Da to 1015 kDa.

##### Bottlebrush Polymer Library Backbone Synthesis and Oligonucleotide Synthesis

Solid-phase synthesis was carried out on model Dr. Oligo 48 (Applied Biosystems, Inc., Foster City, CA). All natural/modified DNA/RNA phosphoramidites were purchased from Glen Research Co. (Sterling, VA, USA). Modified phosphoramidites for the bottlebrush polymer backbone were synthesized as described. Polymer backbones containing Fmoc group were deprotected on column in DMF with 20% piperidine 3× and then washed with DMF 2×. All bottlebrush polymer backbone and oligonucleotides were cleaved from the CPG support and deprotected in aqueous ammonium hydroxide solution (28-30% $NH_3$) at room temperature for 24 h. All digital bottlebrush polymer backbones and oligonucleotide strands were purified by Glen-Pak™ purification column and dried *in vacuo*.

**Cell Culture and Animals**

NCI-H358 cells were cultured in RPMI 1640 media supplemented with 10% fetal bovine serum (FBS) and 1% antibiotics (penicillin and streptomycin). GM03989 and HEP-G2 cells were cultured in EMEM 1640 media supplemented with 10% FBS and 1% antibiotics. All cells were cultured at 37 °C in a humidified atmosphere containing 5% $CO_2$. C57BL/6 and DBA/1J mice (6-8 weeks old) were purchased from Charles River Laboratory (Wilmington, MA, USA). Animals were housed at Northeastern University DLAM animal facilities. Animal protocols (protocol number: 22-0309R) were approved by the Institutional Animal Care and Use Committee of Northeastern University and carried out in accordance with the approved guidelines.

**Cellular Uptake**

All cells were cultured in a 24-well plate with a cell seeding density of $1.0\times10^6$ cells per well in 1 mL complete medium for 24 h at 37 °C. Next, cells were washed twice with 1× PBS and treated with fluorescently labeled test articles and controls, which were dissolved in serum-free culture medium at 0.5 μM concentrations. Cells were further incubated with the samples for 4 h at 37 °C, before being washed with PBS and treated with 100 μL of 0.25% trypsin/EDTA solution (Gibco, USA) followed by 700 μL of cold PBS. Detached cells were collected for flow cytometry analysis (CytoFLEX Flow Cytometer, Beckman Coulter, Brea, CA, USA).

**Confocal Microscopy**

Cells were cultured in a 24-well glass bottom plate at $1.0\times10^5$ cells per well in 1 mL complete RPMI medium for 24 h at 37 °C. On the following day, cells were washed with PBS 3×, and treated with fluorescently labeled test articles in serum-free culture medium at a concentration of 0.5 μM. Cells were further incubated at 37 °C for 8 h. Next, cells were washed with PBS 3× and fixed with 4% paraformaldehyde for 30 min. Finally, the cells were stained with 4′,6-diamidino-2-phenylindole (DAPI) for 10 min before imaged on an LSM-880 confocal laser scanning microscope (Carl Zeiss Ltd., Cambridge, UK). Imaging settings were kept identical.

**Molecular Dynamics Simulation (MD)**

MD simulations were performed using the OPLS4 force field within the Schrödinger Maestro 2024-2 software. The initial polymer structures were constructed using PyMOL 2.5 and subjected to energy minimization pretreatment with the LigPrep tool. At the system construction stage, a polymer-water system was built within a periodic cubic box with dimensions of $8\times8\times8$ $nm^3$. The solvent was modeled using the TIP3P model, and the ionic concentration was set to 0.19 M (NaCl). The initial equilibration process consisted of two stages. First, a 200 ns simulation was carried out in the NVT ensemble, with the temperature controlled by a Nose-Hoover thermostat at 313 K. Subsequently, a 200 ns simulation was performed in the NPT ensemble, and the pressure was regulated using the Martyna-Tobias-Klein method at 1 bar. To investigate the interactions between the polymers and the cell membrane, a DPPC bilayer membrane system was constructed. The pre-equilibrated polymer structures were positioned approximately 1.5 nm away from the membrane surface, forming a composite system of polymer-membrane-water. The interaction simulations were conducted for 1000 ns in the NPγT ensemble at a temperature of 313 K, a pressure of 1 bar, and a surface tension of 0 mN/m. $Na^+$ and $Cl^-$ ions were added to all simulation systems to achieve a physiological salt concentration of 0.19 M and to maintain electrical neutrality. During the simulations, a 9 Å cutoff was applied for nonbonding interactions, and the Particle Mesh Ewald (PME) method was used to handle electrostatic interactions. All hydrogen atoms in covalent bonds were constrained using the SHAKE algorithm, and the integration time step was set to 2 fs. Trajectory data were saved every 5 ps. Each system was processed through the standard Desmond relaxation procedure before the production simulations. The MDAnalysis python package was used for post simulation analysis.

**Plasma Pharmacokinetics (PK)**

Immunocompetent C57BL/6 mice were used to examine the plasma PK. Mice were randomly divided into seven groups. Cy5-labeled samples were i.v. administrated via the tail vein at an equal dosage (0.5 μmol/kg). Blood samples (25 μL) were collected from the submandibular vein at varying time points (24 h, 48 h and 72 h) using BD Vacutainer blood collection tubes with lithium heparin. Heparinized plasma was obtained by centrifugation at 3000 rpm for 20 min, aliquoted into a 96-well plate, and measured on a Synergy™ Neo2 Multi-Mode microplate reader (BioTek Instruments Inc., Winooski, VT, USA). Standard curves were established for each sample, where samples of known quantities were incubated with freshly collected plasma for 1 h at room temperature before fluorescence was measured.

**Multiplex Analysis of Cytokines**

The multiplexing analysis was performed using the Luminex™ 200 system (Luminex, Austin, TX, USA) by Eve Technologies Corp. (Calgary, Alberta, Canada). Ten markers were simultaneously measured in the samples using Mouse Cytokine 10-Plex Discovery Assay® (MilliporeSigma, Burlington, MA, USA) according to the manufacturer's protocol.

**Biodistribution**

C57BL/6 mice (8-10 weeks old, female) were injected i.v. with Cy5-labeled test articles and controls at a dosage of (0.5 μmol/kg). Major organs and tissues were collected and weighed 72 h post injection. Organs and tissues were cut and homogenized in T-PER™ Tissue Protein Extraction Reagent (Thermo Fisher, Waltham, MA, USA) supported with 0.5% Triton X-100 using a BeadBlaster D2400-R refrigerated homogenizer (Benchmark scientific, Sayreville, NJ, USA). After centrifugation, the supernatants were aliquoted into a 96-well plate and measured for fluorescence intensity on a Synergy™ Neo2 Multi-Mode microplate reader (BioTek Instruments Inc., Winooski, VT, USA). The amounts of samples in the supernatant were estimated using standard curves established for each sample.

**Collagen Induced Arthritis Animal Model**

Type II collagen (CII, Sigma-Aldrich Co., St. Louis, MO, USA) and Complete Freund adjuvant (CFA, InvivoGen Inc., San Diego, CA, USA) were used to prepare emulsions at a 1:1 volume ratio using a BeadBlaster D2400-R refrigerated homogenizer (Benchmark scientific, Sayreville, NJ, USA). The emulsion was visually inspected to confirm uniformity and density, which resembles thick whipped cream. The emulsion was then transferred into 1 mL immunization syringes, which were kept on ice until injection. On Day 0, DBA/1J mice (8 weeks; female) were anesthetized with isoflurane prior to subcutaneous administration of 50 μL emulsion (100 μg CII) at a site approximately 1.5 cm distal to the base of the tail. Immunized mice were randomized and assigned to experimental groups and controls. On Day 7, mice were administered in the tail vein with Q16-ASO or controls (13 mg/kg, approximately 20 nmol/animal, ASO basis). On Day 14, mice received a booster subcutaneous immunization of the same dosage as the first injection on the opposite side of the tail. Mice were monitored daily for signs of arthritis, injection site integrity, weight, and signs of discomfort.

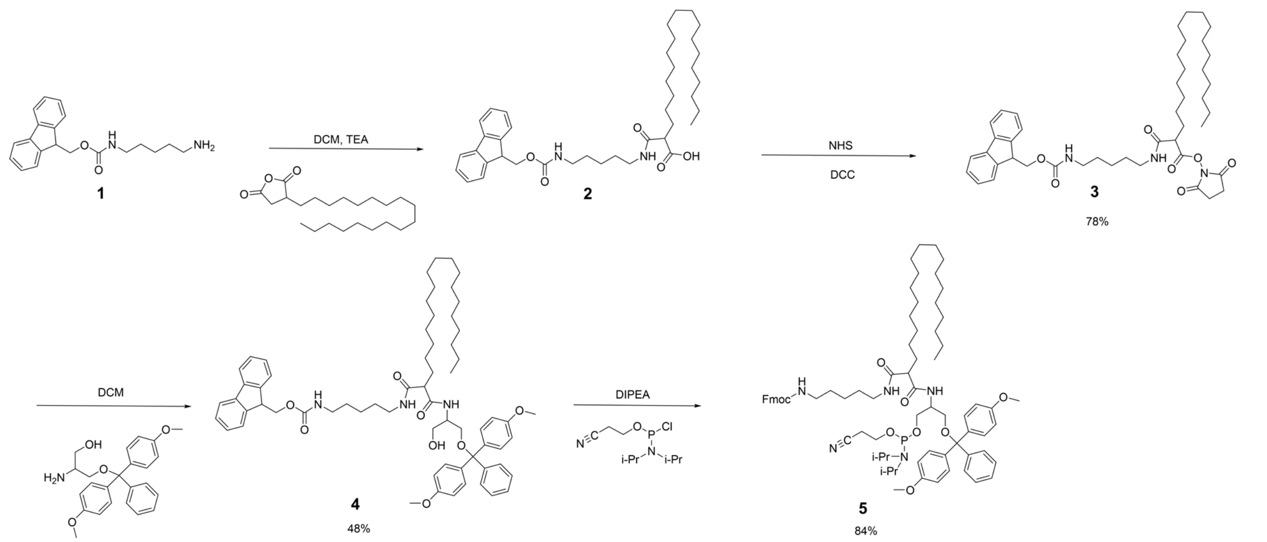

**Scheme S1.** Synthesis of C18 phosphoramidite (**5**)

**Synthesis of C18-NHS (3)**

Octadecylsuccinic anhydride (1.58 g, 5.00 mmol), and (9H-fluoren-9-yl)methyl (5-aminopentyl)carbamate (1.73 g, 5.00 mmol) were dissolved in anhydrous dichloromethane (DCM, 50 mL) under nitrogen atmosphere. Triethylamine (TEA, 0.70 mL, 5.00 mmol) was added, and the reaction mixture was stirred at room temperature overnight. Thereafter, *N*-hydroxysuccinimide (NHS, 0.63 g, 5.50 mmol) and dicyclohexylcarbodiimide (DCC, 1.13 g, 5.50 mmol) were added to the mixture. A white precipitate was observed and removed by filtration. The filtrate was concentrated under reduced pressure, and the residue was purified by column chromatography (30%-60% ethyl acetate in hexane) to give C18-NHS (**3**) as a white solid (2.63 g, 78% yield).

**$^1$H NMR (700 MHz, $CDCl_3$)** δ 7.69 (d, J = 7.5 Hz, 2H), 7.52 (d, J = 7.5 Hz, 2H), 7.33 (t, J = 7.4 Hz, 2H), 7.24 (t, J = 7.4 Hz, 2H), 4.84 (dt, J = 23.3, 6.0 Hz, 1H), 4.33 (d, J = 7.0 Hz, 2H), 4.14 (t, J = 7.0 Hz, 1H), 4.12 – 4.00 (m, 2H), 3.15 – 3.06 (m, 2H), 2.78 – 2.64 (m, 5H), 1.97 (s, 1H), 1.72 (dddd, J = 15.8, 10.4, 5.9, 3.6 Hz, 1H), 1.66 – 1.50 (m, 3H), 1.46 (p, J = 7.2 Hz, 2H), 1.41 – 1.33 (m, 1H), 1.33 – 1.28 (m, 1H), 1.26 – 1.18 (m, 3H), 1.20 – 1.16 (m, 23H), 1.04 (s, 1H), 0.81 (t, J =

7.1 Hz, 3H). **$^{13}$C NMR (176 MHz, $CDCl_3$)** δ 169.79, 169.42, 167.92, 155.40, 142.99, 140.29, 126.64, 126.00, 124.02, 118.94, 65.43, 63.85, 63.75, 59.38, 46.30, 40.06, 39.81, 38.09, 34.87, 32.84, 31.65, 30.92, 30.91, 30.78, 30.22, 28.69, 28.66, 28.64, 28.61, 28.53, 28.47, 28.45, 28.39, 28.35, 28.33, 28.29, 27.15, 27.13, 25.77, 25.55, 24.56, 24.54, 23.87, 22.07, 21.68, 20.04, 13.18, 13.11.

**Synthesis of C18-DMT-OH (4)**

C18-NHS (**3**) (1.01 g, 3.00 mmol) was dissolved in anhydrous DCM (10 mL). Serinol-DMT (1.88 g, 4.00 mmol) was dissolved separately in DCM (5 mL) and added dropwise to a stirring solution of C18-NHS. The reaction mixture was allowed to stir overnight at room temperature. The reaction was monitored by thin-layer chromatography (TLC) using ethyl acetate:hexane (50:50, v:v), where the product showed an Rf of 0.4. Upon completion, the mixture was concentrated under reduced pressure, and the crude product was purified by column chromatography (20%-60% ethyl acetate in hexane) to afford C18-DMT-OH (**4**) as a pale yellow solid (0.87 g, 48% yield).

**$^{1}$H NMR (700 MHz, $CDCl_3$)** δ 7.68 (t, J = 6.0 Hz, 1H), 7.53 – 7.49 (m, 1H), 7.34 – 7.29 (m, 1H), 7.25 – 7.18 (m, 3H), 6.75 (t, J = 8.5 Hz, 1H), 4.31 (dt, J = 7.0, 2.2 Hz, 0H), 3.70 (d, J = 1.9 Hz, 2H), 1.54 (q, J = 7.5 Hz, 1H), 1.17 (d, J = 8.3 Hz, 10H), 1.13 (s, 1H), 0.81 (t, J = 7.0 Hz, 1H). **$^{13}$C NMR (176 MHz, $CDCl_3$)** δ 174.25, 172.09, 157.56, 157.53, 155.49, 143.50, 143.00, 142.98, 140.28, 134.67, 134.56, 128.94, 128.91, 126.99, 126.94, 126.93, 126.90, 126.62, 125.99, 125.95, 125.90, 124.05, 124.02, 118.92, 112.27, 112.23, 112.19, 65.46, 65.40, 63.53, 63.28, 63.07, 62.72, 62.18, 62.05, 54.20, 54.17, 50.54, 50.32, 46.27, 42.36, 40.69, 39.76, 39.73, 37.18, 37.12, 35.92, 31.51, 31.24, 30.91, 30.22, 28.69, 28.68, 28.64, 28.63, 28.59, 28.48, 28.47, 28.44, 28.41, 28.35, 28.33, 27.21, 27.15, 26.33, 26.04, 22.09, 22.00, 21.67, 13.11.

**Synthesis of C18 phosphoramidite (5)**

C18-DMT-OH (**5**) (1.74 g, 4.00 mmol) was placed in a flame-dried flask under nitrogen atmosphere and dissolved in anhydrous DCM (12 mL) containing *N,N*-diisopropylethylamine (DIPEA, 3.50 mL). The solution was chilled in an ice bath. 2-Cyanoethyl *N,N*-diisopropylchlorophosphoramidite (1.90 g, 8.00 mmol) was dissolved in DCM (4 mL) and added dropwise to the stirring reaction mixture. The solution was stirred vigorously at 0 °C for 20 min, then allowed to warm to room temperature and stirred for an additional 40 min. Upon completion, excess ethyl acetate (3 mL) was added to quench the reaction. The mixture was washed with saturated sodium bicarbonate solution (10 mL), and the organic phase was separated, dried over anhydrous sodium sulfate, and filtered. The filtrate was concentrated under reduced pressure, and the crude product was purified by column chromatography using a mixture of hexane:ethyl acetate:TEA (67:33:1, v:v:v) to afford C18 phosphoramidite (**5**) as a yellow oil (4.20 g, 84% yield).

**$^{1}$H NMR (700 MHz, $CDCl_3$)** δ 7.71 – 7.65 (m, 3H), 7.52 (dd, J = 8.0, 4.7 Hz, 3H), 7.36 – 7.28 (m, 6H), 7.26 – 7.17 (m, 10H), 7.16 – 7.10 (m, 2H), 6.74 (dp, J = 9.3, 2.8 Hz, 6H), 4.31 (dd, J = 11.2, 7.2 Hz, 2H), 4.01 – 3.92 (m, 2H), 3.70 (s, 8H), 3.64 – 3.52 (m, 2H), 3.46 (dqd, J = 13.6, 6.6, 3.3 Hz, 2H), 3.16 – 3.00 (m, 3H), 2.67 – 2.38 (m, 7H), 1.59 – 1.53 (m, 2H), 1.42 – 1.24 (m, 3H), 1.17 (dt, J = 11.9, 4.5 Hz, 46H), 1.13 (d, J = 6.8 Hz, 5H), 1.12 – 1.04 (m, 9H), 1.06 – 0.98 (m, 11H), 0.84 – 0.75 (m, 6H). **$^{13}$C NMR (176 MHz, $CDCl_3$)** δ 158.46, 156.47, 144.04, 141.32, 135.99, 130.10, 130.07, 130.05, 128.18, 127.84, 127.81, 127.66, 127.03, 126.80, 125.08, 121.02, 119.96, 119.95, 113.13, 113.10, 86.00, 66.50, 64.32, 64.22, 58.42, 55.22, 55.21, 55.19, 49.72, 49.68, 47.31, 46.16, 43.13, 43.11, 43.06, 41.64, 40.84, 31.95, 31.25, 29.73, 29.68, 29.64, 29.57, 29.53, 29.50,

29.46, 29.38, 28.24, 27.08, 24.64, 24.62, 24.60, 24.58, 23.14, 23.09, 23.00, 22.71, 20.40, 20.36, 14.22, 14.15.

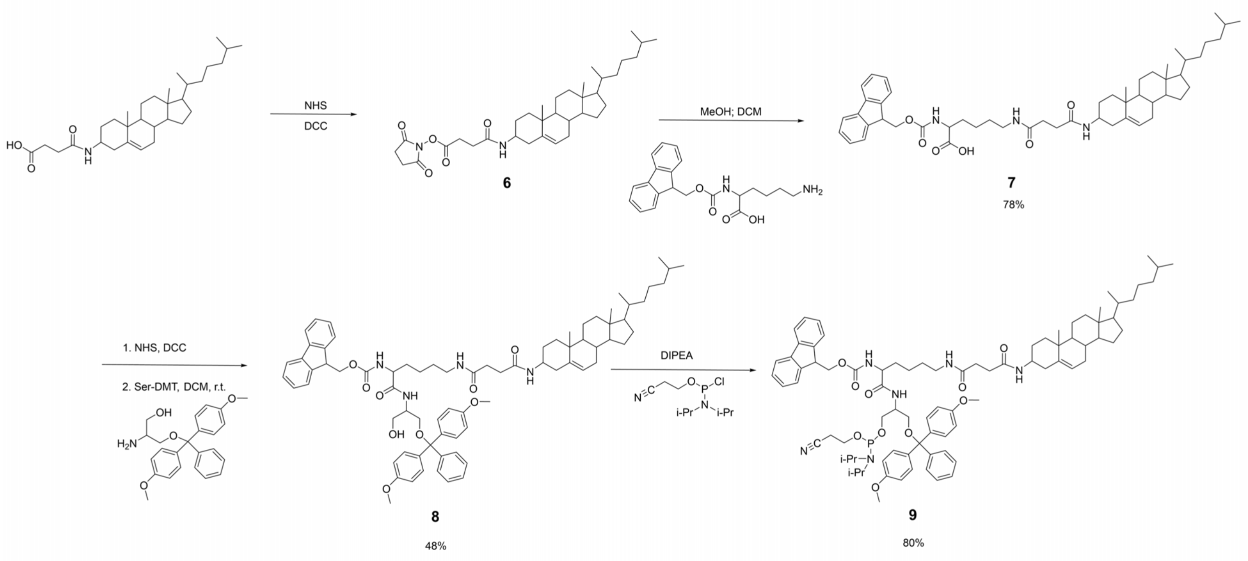


**Scheme S2.** Synthesis of cholesterol phosphoramidite (**9**)

**Synthesis of cholesterol-Lys-OH (7)**

Cholesterylamine (1.93 g, 5.00 mmol) was dissolved in anhydrous DCM (10 mL), followed by the addition of NHS (0.63 g, 5.50 mmol) and DCC (1.13 g, 5.50 mmol). The reaction mixture was stirred at room temperature until a white precipitate formed. The precipitated dicyclohexylurea (DCU) was removed by filtration. To the filtrate, Fmoc-Lys-OH (2.01 g, 5.00 mmol) was added, and the mixture was stirred overnight at room temperature. Thereafter, the reaction mixture was concentrated under reduced pressure, and the residue was purified by column chromatography using 100% ethyl acetate as the eluent to afford cholesterol-Lys-OH (**7**) as a white solid (2.68 g, 78% yield).

**$^1$H NMR (700 MHz, $CDCl_3$)** δ 7.95 (s, 1H), 7.69 (d, J = 7.5 Hz, 2H), 7.54 (t, J = 6.4 Hz, 2H), 7.33 (t, J = 7.4 Hz, 2H), 7.24 (t, J = 7.4 Hz, 2H), 5.96 (t, J = 5.8 Hz, 1H), 5.59 (d, J = 8.2 Hz, 1H), 5.24 (d, J = 5.1 Hz, 1H), 4.67 (q, J = 6.8 Hz, 1H), 4.55 – 4.48 (m, 1H), 4.41 – 4.32 (m, 2H), 4.17 (t, J =

7.1 Hz, 1H), 4.05 (q, J = 7.1 Hz, 1H), 3.22 (hept, J = 6.6 Hz, 2H), 2.89 (s, 2H), 2.81 (s, 2H), 2.78 (s, 5H), 2.74 (s, 1H), 2.63 – 2.52 (m, 3H), 2.38 (t, J = 6.9 Hz, 2H), 2.28 – 2.17 (m, 3H), 1.98 (s, 1H), 1.94 – 1.82 (m, 4H), 1.79 – 1.71 (m, 4H), 1.68 – 1.60 (m, 1H), 1.57 – 1.32 (m, 7H), 1.32 – 1.25 (m, 4H), 1.22 – 1.14 (m, 3H), 1.12 – 0.86 (m, 13H), 0.88 – 0.82 (m, 4H), 0.80 (dd, J = 6.6, 3.3 Hz, 8H), 0.59 (s, 3H). **$^{13}$C NMR (176 MHz, $CDCl_3$)** δ 171.55, 170.87, 170.17, 168.52, 168.00, 167.87, 167.14, 161.57, 154.64, 142.82, 142.61, 140.28, 138.56, 126.71, 126.09, 126.07, 124.10, 121.63, 118.96, 73.85, 73.34, 66.24, 59.39, 58.49, 55.62, 55.09, 51.22, 48.94, 46.09, 41.26, 38.67, 38.49, 37.41, 37.12, 37.01, 36.91, 36.44, 35.91, 35.86, 35.51, 35.50, 35.15, 34.76, 30.87, 30.82, 30.77, 30.64, 30.44, 30.22, 30.05, 28.92, 28.68, 28.03, 27.60, 27.20, 26.99, 26.69, 26.60, 25.32, 24.59, 24.55, 24.43, 24.38, 23.23, 22.80, 21.81, 21.55, 20.44, 20.04, 19.98, 18.26, 17.69, 13.18, 10.82.

**Synthesis of cholesterol-DMT-OH (8)**

Cholesterol-Lys-OH (5.00 mmol) was dissolved in anhydrous DCM (10 mL), followed by the addition of NHS (0.63 g, 5.50 mmol) and DCC (1.13 g, 5.50 mmol). The reaction mixture was stirred at room temperature until a white precipitate formed. The precipitated DCU was removed by filtration. Serinol-DMT (5.5 mmol) was dissolved separately in DCM (5 mL) and added dropwise to the filtered solution. The reaction mixture was allowed to stir overnight at room temperature. Reaction progress was monitored by TLC using ethyl acetate:hexane (50:50, v:v), where the product showed a Rf of 0.6. Upon completion, the mixture was concentrated under reduced pressure, and the crude product was purified by column chromatography (20%-60% ethyl acetate in hexane) to afford cholesterol-DMT-OH (**8**) as a white solid (0.87 g, 48% yield).

**$^{1}$H NMR (700 MHz, $CDCl_3$)** δ 7.68 (d, J = 7.4 Hz, 2H), 7.50 (d, J = 7.5 Hz, 1H), 7.46 (d, J = 7.5 Hz, 1H), 7.31 (td, J = 8.3, 4.7 Hz, 6H), 7.26 – 7.13 (m, 13H), 7.15 – 7.07 (m, 1H), 6.79 – 6.72 (m,

6H), 6.72 (s, 2H), 4.26 (ddd, J = 22.5, 15.5, 8.3 Hz, 2H), 4.08 (s, 2H), 4.11 – 4.04 (m, 1H), 4.06 – 3.99 (m, 1H), 3.77 – 3.69 (m, 5H), 3.67 (dd, J = 11.3, 7.2 Hz, 7H), 3.61 (s, 1H), 3.63 – 3.58 (m, 1H), 3.26 – 3.20 (m, 1H), 3.19 (s, 1H), 3.22 – 3.14 (m, 2H), 3.11 (q, J = 6.3 Hz, 1H), 2.47 – 2.34 (m, 2H), 1.97 (d, J = 2.0 Hz, 2H), 1.95 – 1.88 (m, 1H), 1.86 (s, 1H), 1.82 – 1.70 (m, 2H), 1.67 – 1.60 (m, 1H), 1.48 – 1.39 (m, 3H), 1.38 – 1.23 (m, 3H), 1.29 (s, 5H), 1.19 (td, J = 7.1, 2.2 Hz, 3H), 1.12 – 1.05 (m, 2H), 1.05 – 0.96 (m, 3H), 0.96 – 0.82 (m, 4H), 0.79 (dq, J = 5.4, 2.2 Hz, 4H), 0.61 – 0.56 (m, 2H). **$^{13}$C NMR (176 MHz, $CDCl_3$)** δ 170.16, 157.62, 157.55, 157.53, 143.50, 143.47, 142.85, 140.25, 139.12, 134.58, 128.93, 128.90, 127.71, 127.00, 126.98, 126.95, 126.72, 126.07, 126.03, 125.94, 124.09, 119.98, 118.96, 118.94, 118.72, 112.28, 112.23, 85.44, 85.38, 66.04, 63.14, 62.76, 62.51, 62.31, 62.26, 62.03, 59.39, 58.50, 55.67, 55.10, 54.21, 54.18, 54.16, 53.82, 50.54, 50.29, 50.27, 49.05, 48.78, 46.06, 41.27, 38.71, 38.49, 38.15, 37.71, 37.62, 37.12, 36.78, 35.50, 35.47, 35.16, 34.78, 30.98, 30.92, 30.80, 30.22, 28.69, 28.08, 28.00, 27.90, 27.21, 26.99, 23.25, 22.80, 22.36, 22.25, 21.81, 21.55, 21.34, 21.11, 20.05, 19.93, 18.30, 18.26, 17.69, 13.18, 10.83.

**Synthesis of cholesterol phosphoramidite (9)**

Cholesterol-DMT-OH (2.06 g, 4.00 mmol) was placed in a flame-dried flask under nitrogen atmosphere and dissolved in anhydrous DCM (12 mL) containing DIPEA (3.50 mL). The solution was chilled in an ice bath. 2-Cyanoethyl *N,N*-diisopropylchlorophosphoramidite (1.90 g, 8.00 mmol) was dissolved in DCM (4 mL) and added dropwise to the stirred reaction mixture. The solution was stirred vigorously at 0 °C for 20 min, then allowed to warm to room temperature and stirred for an additional 40 min. Upon completion, excess ethyl acetate (3 mL) was added to quench the reaction. The mixture was washed with saturated sodium bicarbonate solution (10 mL), and the organic phase was separated, dried over anhydrous sodium sulfate, and filtered. The filtrate was concentrated under reduced pressure, and the crude product was purified by column chromatography using a mixture of hexane:ethyl acetate: TEA (67:33:1, v:v:v) to afford cholesterol phosphoramidite (**9**) as a white solid (4.50 g, 80% yield).

**$^1$H NMR (700 MHz, $CDCl_3$)** δ 7.69 (q, J = 3.6 Hz, 2H), 7.47 (q, J = 8.2 Hz, 0H), 7.36 – 7.28 (m, 4H), 7.23 (s, 1H), 7.22 (d, J = 3.5 Hz, 2H), 7.13 – 7.09 (m, 2H), 6.78 – 6.70 (m, 4H), 6.61 (s, 1H), 4.25 (s, 2H), 4.14 (s, 2H), 4.11 – 4.02 (m, 1H), 3.82 – 3.78 (m, 1H), 3.76 (s, 1H), 3.74 – 3.64 (m, 6H), 3.45 (d, J = 12.9 Hz, 1H), 3.44 (s, 3H), 3.23 (s, 2H), 3.10 (s, 1H), 2.57 (s, 1H), 2.48 – 2.40 (m, 1H), 2.16 (s, 2H), 1.84 (d, J = 10.2 Hz, 1H), 1.37 (d, J = 11.1 Hz, 1H), 1.34 (s, 2H), 1.25 – 1.12 (m, 11H), 1.12 – 1.05 (m, 10H), 1.04 (s, 6H), 1.07 – 0.99 (m, 3H), 1.01 (s, 2H), 0.85 – 0.75 (m, 3H). **$^{13}$C NMR (176 MHz, $CDCl_3$)** δ 158.45, 141.26, 130.10, 128.16, 127.87 – 125.41 (m), 125.25, 119.95, 113.11, 59.53, 55.18, 47.12, 45.01, 43.07 (d, J = 14.1 Hz), 38.16, 31.25, 24.60, 22.71, 20.38, 14.14, 1.03.

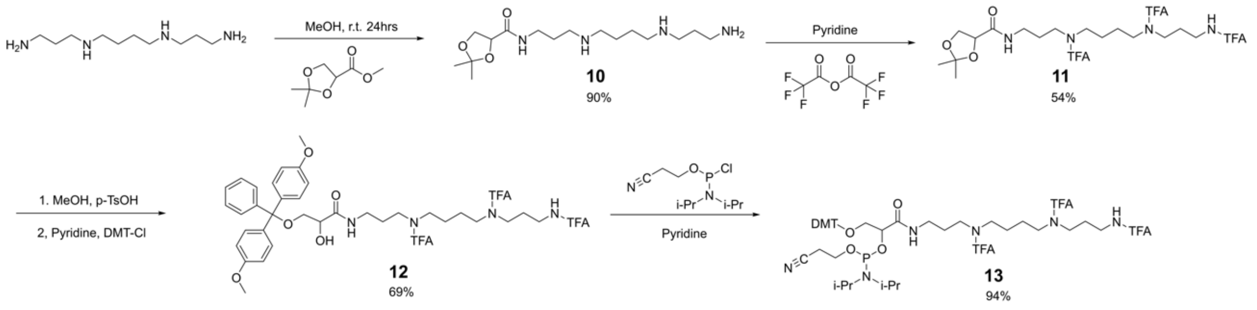


**Scheme S3.** Synthesis of spermine phosphoramidite (**13**)

**Synthesis of spermine-prot-diol (10)**

Methyl (S)-2,2-dimethyl-1,3-dioxalane-4-carboxylate (0.45 mL, 3 mmol) and spermine (2.40 g, 12 mmol) were dissolved in anhydrous MeOH (1 mL) and the resulting solution was stirred at room temperature for 48 h. The solvent was evaporated under reduced pressure, and the residue was purified by column chromatography using a mixture of MeOH:$MeNH_2$:$H_2O$ (90:6:4, v:v:v) to give spermine-prot-diol (**10**) as yellow oil (0.90 g, 90% yield).

**$^1$H NMR (700 MHz, $CDCl_3$)** δ 7.22 (dt, J = 20.5, 5.8 Hz, 1H), 4.40 (dd, J = 7.6, 5.4 Hz, 1H), 4.21 (t, J = 8.2 Hz, 1H), 4.00 (dd, J = 8.8, 5.4 Hz, 1H), 3.34 – 3.23 (m, 2H), 2.69 (t, J = 6.9 Hz, 1H), 2.60 (td, J = 6.9, 3.4 Hz, 3H), 2.56 – 2.50 (m, 4H), 2.50 – 2.41 (m, 3H), 1.62 (p, J = 6.6 Hz, 2H), 1.57 (q, J = 7.0 Hz, 1H), 1.49 – 1.43 (m, 3H), 1.41 (s, 3H), 1.32 (s, 3H).

**Synthesis of spermine-prot-diol-TFA (11)**

Spermine-prot-diol (**10**) (0.20 g, 0.60 mmol) was co-evaporated with anhydrous pyridine (5 mL) 3x and redissolved in pyridine (6 mL). Trifluoroacetic anhydride (0.50 mL, 3.60 mmol) was added dropwise to a cooled and stirring solution at 0 °C. The solution was allowed to warm to room temperature under stirring. The reaction was quenched with sodium bicarbonate and extracted with DCM. The combined organic extracts were washed with brine, dried over anhydrous sodium sulfate, and concentrated under vacuum. The residual oil was purified by column chromatography with DCM:MeOH (98:2, v:v) as the eluent to give spermine-prot-diol-TFA (**11**) as a white foam (0.20 g, 54% yield).

**$^1$H NMR (700 MHz, $CDCl_3$)** δ 4.10 (dq, J = 12.4, 4.3 Hz, 1H), 4.05 (q, J = 7.1 Hz, 4H), 3.89 – 3.74 (m, 1H), 3.42 (q, J = 6.4 Hz, 1H), 3.35 (q, J = 7.7 Hz, 5H), 3.32 – 3.18 (m, 2H), 1.84 – 1.76 (m, 1H), 1.56 (dt, J = 7.7, 3.9 Hz, 2H), 1.35 (d, J = 10.1 Hz, 1H), 1.19 (t, J = 7.1 Hz, 7H), 1.04 (s, 2H), 0.77 (s, 1H). **$^{13}$C NMR (176 MHz, $CDCl_3$)** δ 170.23, 70.93, 59.41, 30.22, 24.71, 20.05, 13.18.

**Synthesis of spermine-DMT (12)**

To a solution of spermine-prot-diol-TFA (**11**) (0.50 g, 0.75 mmol) in MeOH (4 mL), p-toluenosulfonic acid (0.30 g, 1.50 mmol) was added at room temperature. The mixture was stirred for 30 h at room temperature, and the solvent was evaporated. The mixture was diluted with ethyl acetate and washed with saturated sodium bicarbonate solution (10 mL). The combined organic extracts were dried over anhydrous sodium sulfate and concentrated under vacuum. The resulting crude was co-evaporated with pyridine (5 mL) 4x and dissolved in pyridine (2.50 mL). 4,4-Dimethoxytrityl chloride (0.30 g, 0.98 mmol) was added in portions at room temperature. The mixture was stirred for 3 h at room temperature then quenched with MeOH (1 mL) and saturated sodium bicarbonate solution (10 mL). The mixture was extracted with DCM. The resulting orange oil was purified by chromatography with 2%-10% MeOH in $CH_2Cl_2$ (with 1% of TEA by vol.) as the eluent to give spermine-DMT (**12**) as a white solid (0.49 g, 69% yield).

**$^1$H NMR (700 MHz, $CDCl_3$)** δ 7.47 – 7.39 (m, 2H), 7.30 (dd, J = 7.7, 5.2 Hz, 5H), 7.24 (q, J = 7.3 Hz, 1H), 7.21 – 7.14 (m, 1H), 6.85 (dd, J = 8.9, 3.4 Hz, 3H), 4.21 – 4.11 (m, 1H), 3.81 (s, 4H), 3.55 – 3.26 (m, 6H), 2.07 (s, 1H), 1.95 – 1.84 (m, 2H), 1.84 (d, J = 6.3 Hz, 1H), 1.59 (dp, J = 11.3, 3.9 Hz, 3H), 1.28 (t, J = 7.1 Hz, 1H), 1.13 (s, 1H), 0.09 (s, 1H). **$^{13}$C NMR (176 MHz, $CDCl_3$)** δ 157.62, 146.29, 138.43, 128.93, 128.89, 128.11, 127.00, 126.96, 126.92, 126.84, 126.74, 126.10, 126.07, 112.30, 112.26, 112.15, 85.82, 70.93, 69.96, 63.03, 59.39, 58.50, 54.24, 54.22, 45.97, 43.10, 42.87, 42.75, 37.12, 35.58, 35.51, 35.35, 35.30, 34.98, 30.22, 28.68, 28.16, 26.17, 25.58, 25.50, 24.70, 22.90, 22.78, 20.05, 13.18.

**Synthesis of spermine phosphoramidite (13)**

Spermine-DMT (**12**) (4.10 g, 4 mmol) was placed in a flame-dried flask under nitrogen atmosphere and dissolved in anhydrous DCM (12 mL) containing DIPEA (3.50 mL). The flask was chilled in an ice bath. 2-Cyanothyl *N,N*-diisopropylchlorophosphoramidite (1.90 g, 8 mmol) was dissolved in DCM (4 mL) and added dropwise to stirred reaction mixture. The reaction mixture was allowed to stir vigorously for 20 min before being warmed to room temperature and then stirred for another 120 min. An excess of ethyl acetate was added to the reaction mixture. The mixture was washed with saturated sodium bicarbonate solution (10 mL). After drying over anhydrous sodium sulfate, the mixture was filtered, and the filtrate was concentrated and purified by column chromatography over a mixture of hexane:ethyl acetate:TEA (67:33:1, v:v:v) to afford spermine phosphoramidite (**13**) as a white solid (4.10 g, 94% yield).

**$^1$H NMR (700 MHz, $CDCl_3$)** δ 7.36 (td, J = 8.6, 5.5 Hz, 1H), 7.26 – 7.09 (m, 5H), 6.76 – 6.70 (m, 3H), 4.06 (dq, J = 14.1, 6.6 Hz, 1H), 3.71 (d, J = 6.8 Hz, 4H), 3.61 – 3.54 (m, 1H), 3.52 – 3.18 (m, 5H), 3.19 (s, 1H), 2.74 – 2.64 (m, 1H), 2.60 – 2.53 (m, 1H), 2.49 – 2.39 (m, 1H), 1.97 (s, 1H), 1.87 – 1.68 (m, 2H), 1.46 (td, J = 19.0, 8.5 Hz, 1H), 1.41 (d, J = 6.5 Hz, 1H), 1.40 – 1.33 (m, 1H), 1.20 (dd, J = 10.1, 6.9 Hz, 7H), 1.17 – 1.11 (m, 4H), 1.06 (dd, J = 14.7, 8.8 Hz, 5H), 0.84 – 0.75 (m, 1H). **$^{13}$C NMR (176 MHz, $CDCl_3$)** δ 170.16, 169.90, 157.57, 157.55, 157.53, 157.51, 157.46, 157.43, 143.76, 143.72, 143.67, 143.61, 134.88, 134.82, 134.73, 134.68, 134.61, 129.16, 129.12, 129.11, 129.06, 127.23, 127.22, 127.19, 127.17, 126.72, 126.70, 126.66, 125.90, 125.83, 125.75, 125.69, 117.50, 117.10, 116.79, 116.75, 116.17, 115.88, 114.03, 112.03, 112.01, 111.97, 84.95, 84.88, 84.85, 72.47, 72.39, 72.31, 64.02, 63.59, 59.39, 58.50, 57.72, 57.65, 57.63, 57.13, 57.10, 57.07, 56.96, 56.79, 56.67, 54.20, 54.17, 46.30, 46.07, 45.47, 45.26, 45.17, 44.80, 44.31, 44.27, 43.61, 43.50, 42.66, 42.63, 42.52, 42.45, 42.37, 42.30, 37.12, 35.96, 35.90, 35.55, 35.52, 35.40, 35.38, 35.36, 35.35, 30.22, 28.68, 28.28, 28.21, 28.08, 27.24, 27.20, 26.31, 25.48, 25.40, 24.75, 24.71, 24.68, 24.64, 24.61, 23.62, 23.60, 23.57, 22.89, 21.96, 21.95, 21.88, 21.86, 20.04, 19.59, 19.56, 19.51, 19.47, 19.42, 19.39, 19.11, 19.07, 13.18, 13.11.

**Synthesis of serinol phosphoramidite (16)**

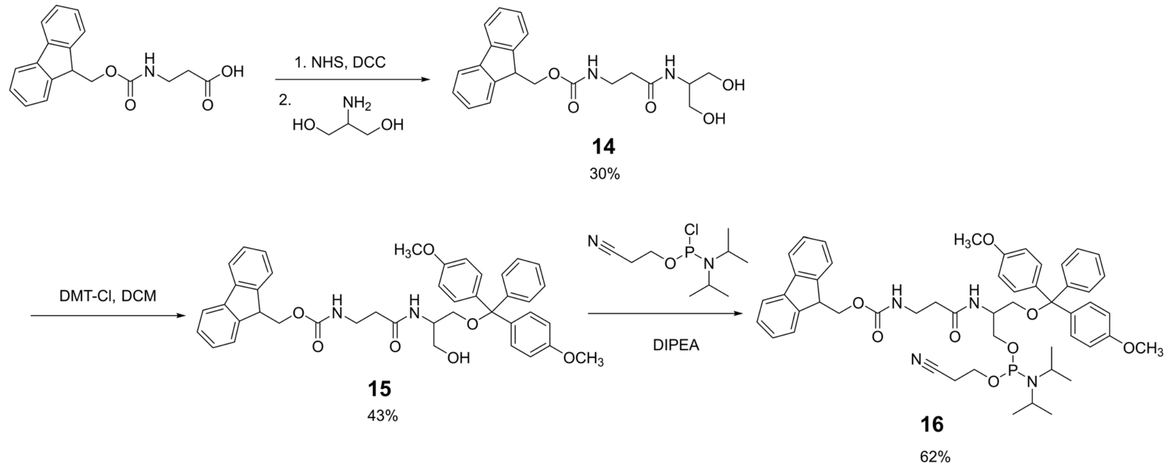


**Scheme S4.** Synthesis of serinol phosphoramidite (**16**)

Fmoc-β-alanine (9.30 g, 30 mmol) was activated with NHS and DCC in a solution of DCM (30 mL) and *N,N*-dimethylformamide (3 mL) and coupled with 2-amino-1,3-propanediol (serinol, 2.70 g, 30 mmol) in pyridine. After overnight stirring, the mixture was co-evaporated with toluene to remove pyridine, and the crude was refluxed in DCM, cooled, filtered, and washed to yield Fmoc-β-Ala-serinol (**14**) as a white solid (3.50 g, 30% yield). This intermediate (10.90 g, 15.30 mmol) was then reacted with 4,4-dimethoxytrityl chloride (5.40 g, 15.90 mmol) in pyridine under nitrogen in an ice bath for 1 h, followed by room temperature stirring overnight. After quenching MeOH and extracted by DCM, the product was purified by column chromatography over silica gel using hexane:ethyl acetate:TEA (90:9:1, v:v:v) to give Fmoc-β-Ala-serinol-DMT (**15**) as a white solid (1.50 g, 43% yield). Finally, Fmoc-β-Ala-serinol-DMT (**15**) (2.80 g, 4 mmol) was reacted with 2-cyanoethyl *N,N*-diisopropylchlorophosphoramidite (1.90 g, 8 mmol) and DIPEA in dry DCM under nitrogen. The product was purified by column chromatography using hexane:ethyl acetate:TEA (60:39:1, v:v:v), to afford serinol phosphoramidite (**16**) as a white solid (2.00 g, 62% yield).

**$^1$H NMR (700 MHz, $CDCl_3$)** δ 7.77 (d, J = 7.5 Hz, 2H), 7.59 (dd, J = 7.6, 4.1 Hz, 2H), 7.46 – 7.37 (m, 4H), 7.32 (d, J = 7.8 Hz, 8H), 7.23 (d, J = 7.4 Hz, 1H), 6.84 (d, J = 8.2 Hz, 4H), 5.84 (d, J = 8.6 Hz, 1H), 5.57 (q, J = 6.7 Hz, 1H), 4.34 (d, J = 7.4 Hz, 2H), 4.16 (dt, J = 29.2, 7.3 Hz, 1H), 3.80 (d, J = 11.5 Hz, 9H), 3.71 – 3.62 (m, 2H), 3.61 – 3.51 (m, 2H), 3.49 (s, 2H), 3.42 – 3.28 (m, 1H), 3.16 (dt, J = 9.3, 6.2 Hz, 1H), 2.60 – 2.47 (m, 2H), 2.38 (dd, J = 13.3, 6.7 Hz, 2H), 1.20 – 1.11 (m, 12H). **$^{13}$C NMR (176 MHz, $CDCl_3$)** δ 170.16, 169.90, 143.76, 143.61, 134.88, 134.82, 134.73, 134.68, 134.61, 129.16, 129.12, 115.88, 84.95, 84.88, 84.85, 72.47, 64.02, 63.59, 59.39, 57.72, 54.17, 46.30, 42.30, 37.12, 35.96, 28.68, 21.86, 13.11.

#### Supplementary Tables

| Sample ID | Backbone Sequence |
|---|---|
| **Q1** | NN NNN NNN NNN NNN NNN NNN NNN NNN NNN N |
| **Q2** | NN NNN NNN NNN NAA AAA ANN NNN NNN NNN N |
| **Q3** | NN NNN NNN AAA NNN NNN NNA AAN NNN NNN N |
| **Q4** | NN NNN NAA NNN NNN AAN NNN NNA ANN NNN N |
| **Q5** | NN NAN NNA NNN NAN NNN ANN NNA NNN ANN N |
| **Q6** | AA ANN NNN NNN NNN NNN NNN NNN NNN NAA A |
| **Q7** | NA NNA NNA NNA NNA NNA NNA NNA NNA NNA N |
| **Q16** | NN NNN NNN NNN NCC CCC CNN NNN NNN NNN N |
| **Q17** | NN NNN NCC NNN NNN CCN NNN NNC CNN NNN N |
| **Q18** | NN NCN NNC NNN NCN NNN CNN NNC NNN CNN N |
| **Q19** | CC CNN NNN NNN NNN NNN NNN NNN NNN NCC C |
| **Q27** | NN NNN NNN NNN NAA SAA ANN NNN NNN NNN N |
| **Q29** | NA NNA NNA NNA NNS NNS NNA NNA NNA NNA N |
| **Q30** | AA ASN NNN NNN NNN NNN NNN NNN NNN SAA A |
| **Q32** | NA NNA NNS NNA NNA NNA NNA NNS NNA NNA N |
| **Q34** | NN NNN NNN NNN NNN ANN NNN NNN NNN NNN N |
| **Q35** | NN NNN NNN NAN NNN NNN NNA NNN NNN NNN N |
| **Q36** | NN NNN NAN NNN NNN ANN NNN NNA NNN NNN N |
| **Q37** | NN NNN ANN NNN ANN NNN ANN NNN ANN NNN N |
| **Q38** | NN NNA NNN NAN NNN ANN NNA NNN NAN NNN N |
| **Q39** | NN NNN NAN NNN NNN SSN NNN NNN NNN NNN N |
| **Q40** | NN NNN NAN NNN NNN SSN NNN NNN ANN NNN N |
| **Q41** | NN NNA NNN NAN NNN SSN NNN NNA NNN NNN N |
| **Q42** | NN NNA NNN NAN NNN SSN NNN ANN NNA NNN N |
| **Q43** | NN NAN NNA NNN NAN SSN ANN NNA NNN ANN N |
| **Q44** | CC CAN NNA NNN NAN SSN ANN NNA NNN ACC C |
| **Q45** | NA NNA NNA NNA NNN SSN NNA NNA NNA NNA N |
| **Q46** | NN NNC NNA NNC NNN NAN NNN CNN ANN CNN N |
| **Q47** | AN NNN NNA NNN NNN ANN CNN NCN NNC NNN C |
| **Q48** | NN NNC NNA NNC NSN NAN NNN CNN ANN CNN N |
| **Q49** | AN NNN NNA NNN SNN ANN CNN NCN NNC NNN C |

**Table S1.** Sequences of polyphosphodiester backbone synthesized in this study. **N**: non-modified (serinol); **A**: C18 alkyl; **C**: cholesterol; **S**: spermine. All monomers (**N**, **A**, **C**, and **S**) are subsequently linked to a polyethylene glycol (PEG, 10 kDa) size chain to afford the bottlebrush architecture.

| Sample ID | Mn (kDa) | Mw (kDa) | PDI | DLS size (nm) |
|---|---|---|---|---|
| **Q1** | 340.91 | 382.71 | 1.12 | 25.0±1.5 |
| **Q2** | 335.76 | 378.92 | 1.13 | 26.0±3.0 |
| **Q3** | 329.56 | 378.15 | 1.15 | 24.0±0.6 |
| **Q4** | 350.14 | 369.77 | 1.06 | 23.0±3.6 |
| **Q5** | 299.47 | 346.93 | 1.16 | 25.0±0.8 |
| **Q6** | 321.80 | 373.53 | 1.16 | 23.4±2.0 |
| **Q7** | 316.58 | 343.75 | 1.09 | 22.9±3.3 |
| **Q16** | 333.10 | 376.65 | 1.13 | 23.6±0.3 |
| **Q17** | 352.54 | 393.21 | 1.12 | 22.2±2.6 |
| **Q18** | 296.92 | 349.83 | 1.18 | 24.8±1.4 |
| **Q19** | 290.98 | 345.93 | 1.19 | 24.3±1.4 |
| **Q27** | 338.71 | 362.59 | 1.07 | 23.7±1.6 |
| **Q29** | 345.13 | 374.31 | 1.08 | 21.9±1.2 |
| **Q31** | 352.33 | 367.11 | 1.04 | 23.0±1.3 |
| **Q32** | 342.01 | 402.29 | 1.18 | 22.9±1.9 |
| **Q34** | 290.44 | 345.63 | 1.19 | 24.8±1.1 |
| **Q35** | 323.56 | 352.16 | 1.09 | 21.8±3.2 |
| **Q36** | 342.13 | 387.21 | 1.13 | 22.4±2.1 |
| **Q37** | 323.19 | 349.12 | 1.08 | 24.3±0.8 |
| **Q38** | 320.98 | 353.23 | 1.10 | 23.1±1.7 |
| **Q39** | 310.09 | 339.75 | 1.10 | 24.8±1.5 |
| **Q40** | 321.06 | 402.42 | 1.25 | 23.6±1.4 |
| **Q41** | 348.91 | 410.29 | 1.18 | 22.6±3.5 |
| **Q42** | 343.09 | 406.73 | 1.19 | 23.3±2.8 |
| **Q43** | 330.19 | 372.32 | 1.13 | 25.3±0.8 |
| **Q44** | 341.02 | 381.03 | 1.12 | 22.7±1.8 |
| **Q45** | 292.47 | 339.92 | 1.16 | 25.2±1.0 |

**Table S2.** Number-average molecular weight ($M_n$), weight-average molecular weight ($M_w$), polydispersity index (PDI), and Z-average hydrodynamic diameter of all bottlebrush polymers tested.

| Sample ID | Half-life (h) | AUC$_{\infty}$ (nmol/mL·h) |
|---|---|---|
| **Q0** | 11.0 | 232.2 |
| **Q1** | 15.8 | 227.7 |
| **Q2** | 8.7 | 208.2 |
| **Q3** | 24.5 | 399.3 |
| **Q4** | 47.4 | 655.9 |
| **Q5** | 19.3 | 295.1 |
| **Q6** | 24.1 | 394.8 |
| **Q7** | 31.9 | 500.4 |
| **Q16** | 11.7 | 250.9 |
| **Q17** | 18.9 | 316.1 |
| **Q18** | 49.0 | 635.5 |
| **Q19** | 18.6 | 302.4 |
| **Q27** | 22.8 | 362.3 |
| **Q29** | 13.5 | 242.8 |
| **Q30** | 43.9 | 571.7 |
| **Q32** | 28.2 | 427.0 |
| **Q34** | 26.0 | 388.5 |
| **Q35** | 35.5 | 511.5 |
| **Q36** | 39.3 | 565.0 |
| **Q37** | 22.3 | 407.4 |
| **Q38** | 22.0 | 343.2 |
| **Q39** | 52.1 | 670.8 |
| **Q40** | 32.5 | 488.5 |
| **Q41** | 38.9 | 559.5 |
| **Q42** | 23.8 | 394.3 |
| **Q43** | 23.6 | 394.8 |
| **Q45** | 25.0 | 419.7 |
| **Q46** | 24.3 | 310.8 |
| **Q47** | 44.9 | 469.0 |
| **Q48** | 14.3 | 224.6 |
| **Q49** | 21.6 | 287.0 |

**Table S3.** Pharmacokinetic parameters. Half-life ($t_{1/2}$) and area under the curve (AUC$_{\infty}$) were obtained by fitting a one-phase exponential decay model.

| Severity score | Degree of inflammation |
|---|---|
| 0 | No visible signs of erythema or swelling |
| 1 | Erythema and mild swelling confined to the tarsals or ankle joint |
| 2 | Erythema and mild swelling extending from the ankle to the tarsals |
| 3 | Erythema and moderate swelling extending from the ankle to the metatarsal joints |
| 4 | Erythema and severe swelling encompassing the ankle, foot, and digits, or presence of ankylosis |

**Table S4.** Arthritis severity evaluation. Arthritis severity was assessed by visually scoring each limb using a standardized 0-4 scale based on the degree of erythema and swelling. Each of the four limbs was scored independently, and the final arthritis severity score for each mouse was calculated as the average of the four limb scores, including both forelimbs and hind limbs. Scoring was performed by three investigators in a blinded manner to ensure consistency across readings.

**Supplementary Figures**

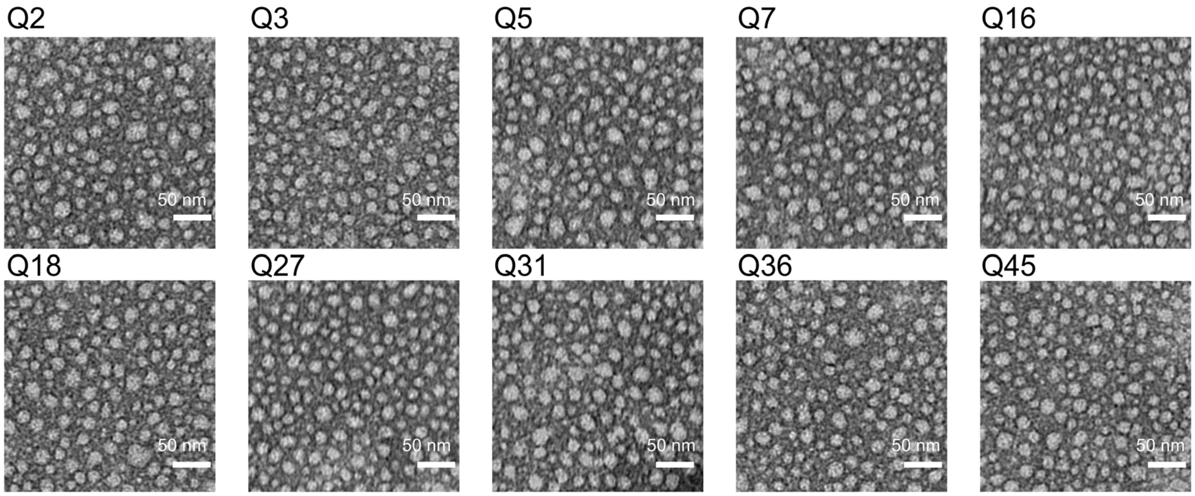


**Figure S1.** Transmission electron microscopy (TEM) images of representative bottlebrush polymers.

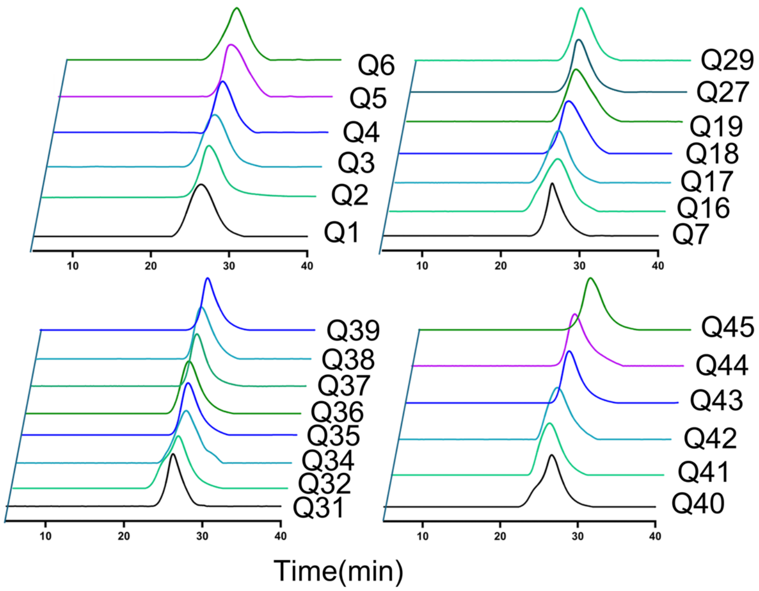


**Figure S2.** Additional size exclusion chromatography (SEC) characterization of bottlebrush polymers.

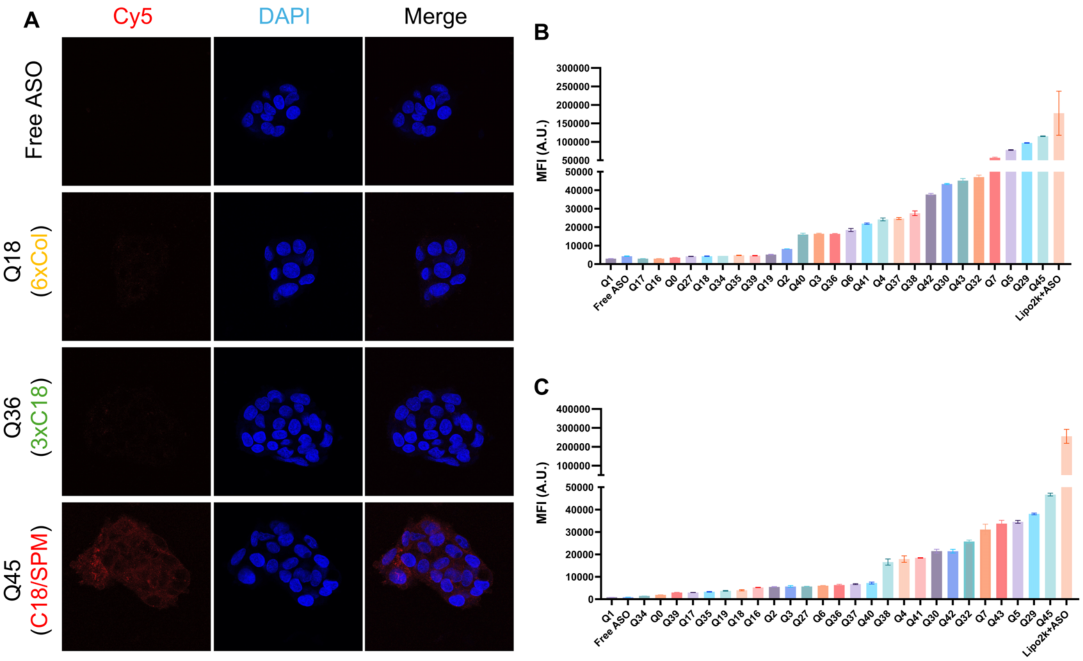


**Figure S3.** Additional confocal microscopy images and cellular uptake results. (A) Confocal microscopy images of HepG2 cells treated with selected Cy5-labeled bottlebrush polymer samples (500 nM) for 8 h (B-C) Mean florescence intensity (MFI) of GM03989 cells (B) and HepG2 cells (C) treated with bottlebrush polymers for 4 h (500 nM) as determined flow cytometry.

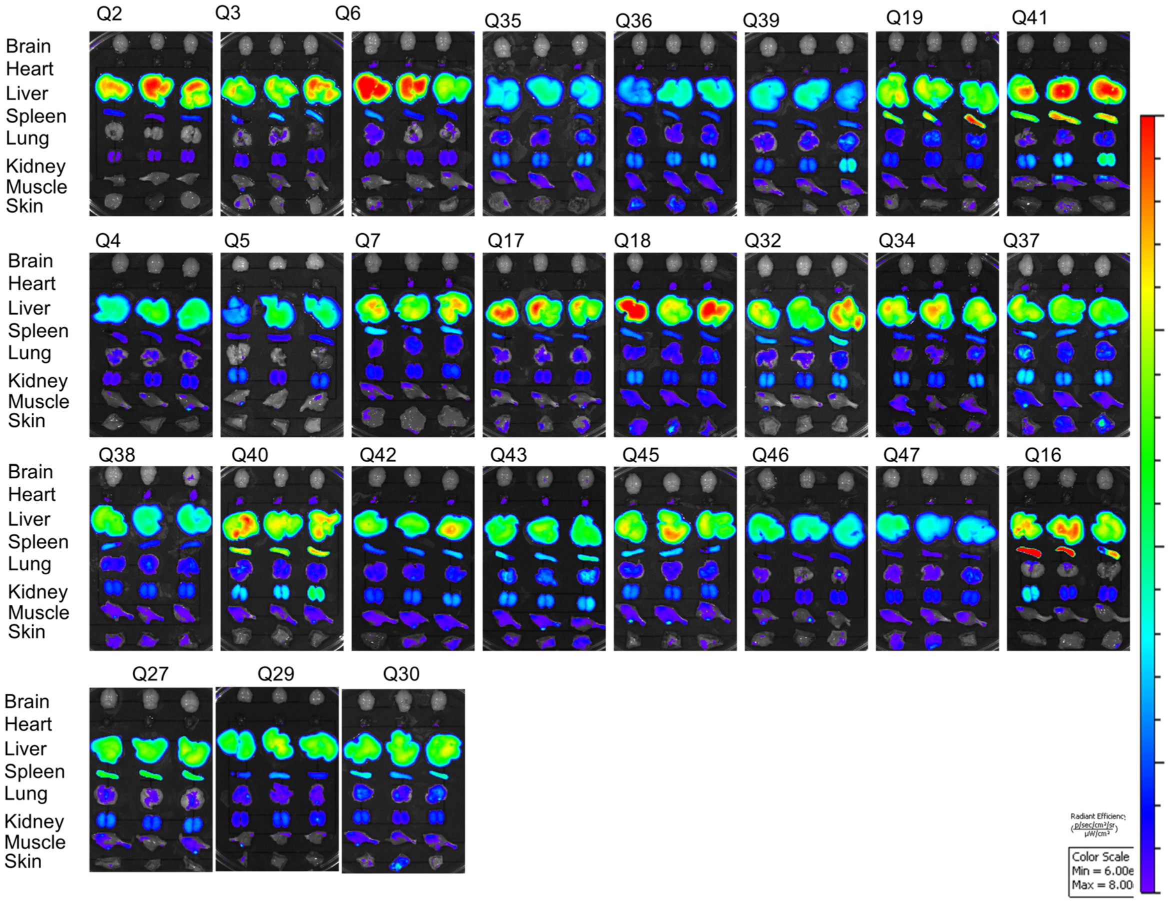


**Figure S4.** *Ex vivo* organ imaging of mice treated with Cy5-labeled digital bottlebrush polymers 72 h after i.v. injection.

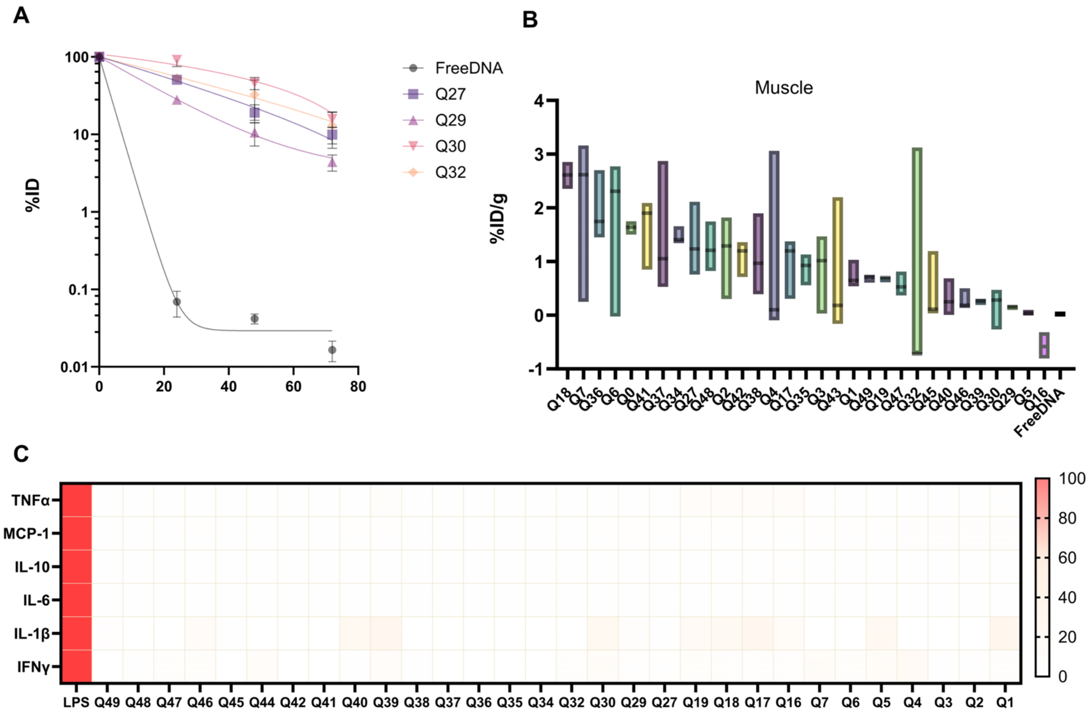


**Figure S5.** Additional plasma pharmacokinetics, muscle biodistribution, and innate immune activation data. (A) Plasma pharmacokinetics of four backbone-modified bottlebrush polymer samples measured over time following intravenous administration. (B) Biodistribution of bottlebrush polymers in muscle (thigh) from high to low. (C) Relative cytokine expression levels (0-100%) in plasma 72 h after i.v. injection of digital bottlebrush polymers.

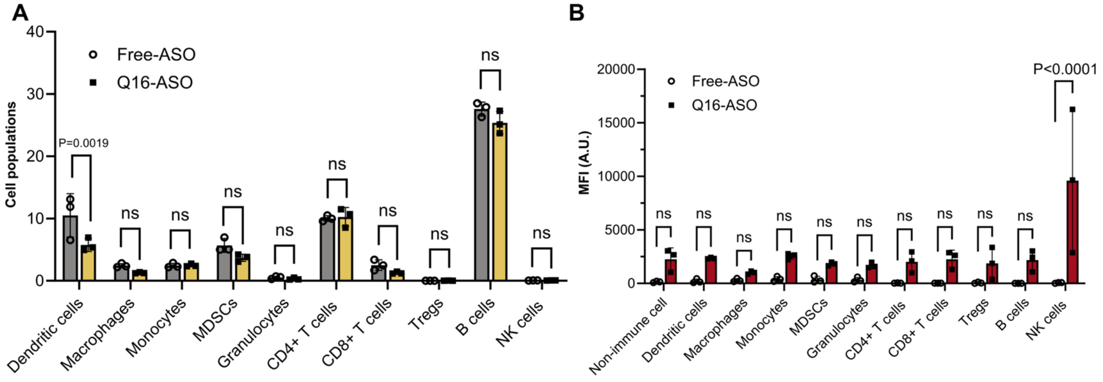


**Fig. S6.** Immune cell frequency and MFI in isolated spleen. (A) Immune cell frequency in spleen 72 h following i.v. administration of free ASO and Cy5-labeled Q16-ASO. (B) Cy5+ immune and non-immune cell MFI in the spleen 72 h following i.v. administration of free ASO and Cy5-labeled Q16-ASO. Statistical analysis was performed using two-way ANOVA.

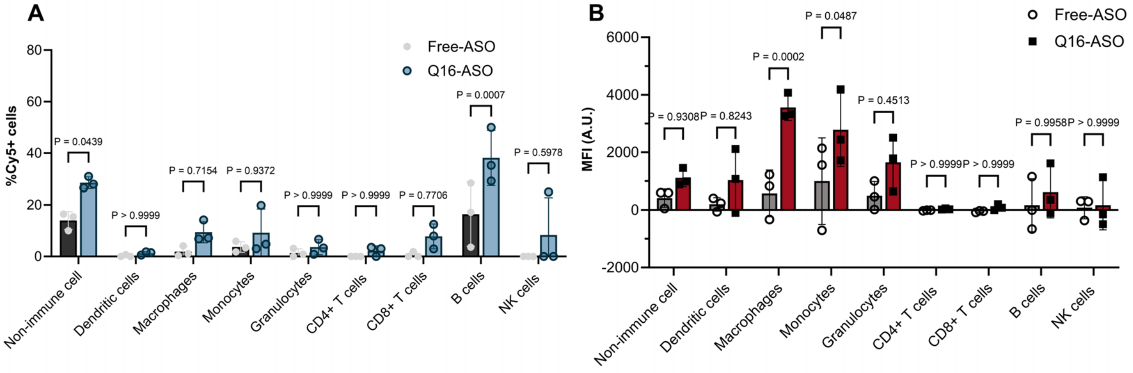


**Figure S7.** Cy5$^+$ immune and non-immune cell populations and MFI in popliteal fossa synovium. (A) Cy5$^+$ cell populations (B) MFI in popliteal fossa synovium 72 h following i.v. administration of Cy5-labeled Q16-ASO. Statistical analysis was performed using two-way ANOVA.

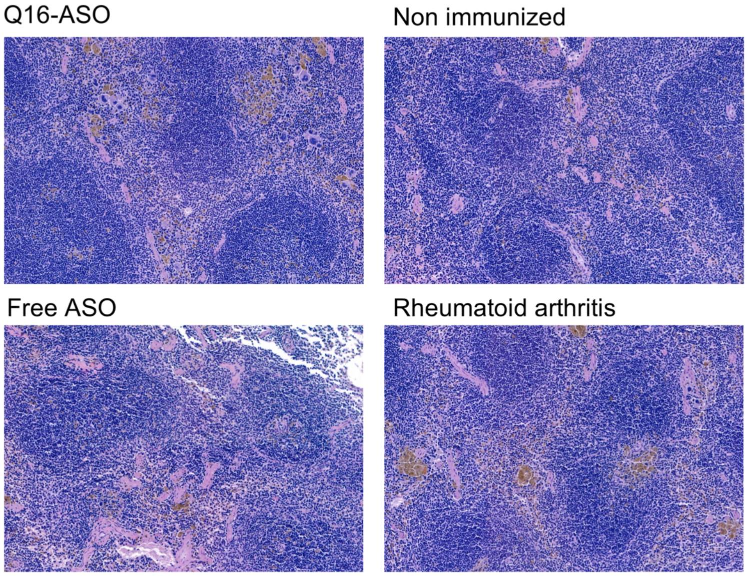


**Figure S8.** H&E staining of spleen sections from Q16-ASO treated, non-immunized (healthy), free ASO-treated, and rheumatoid arthritis mice.

#### Compound 3

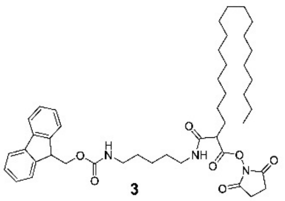

3

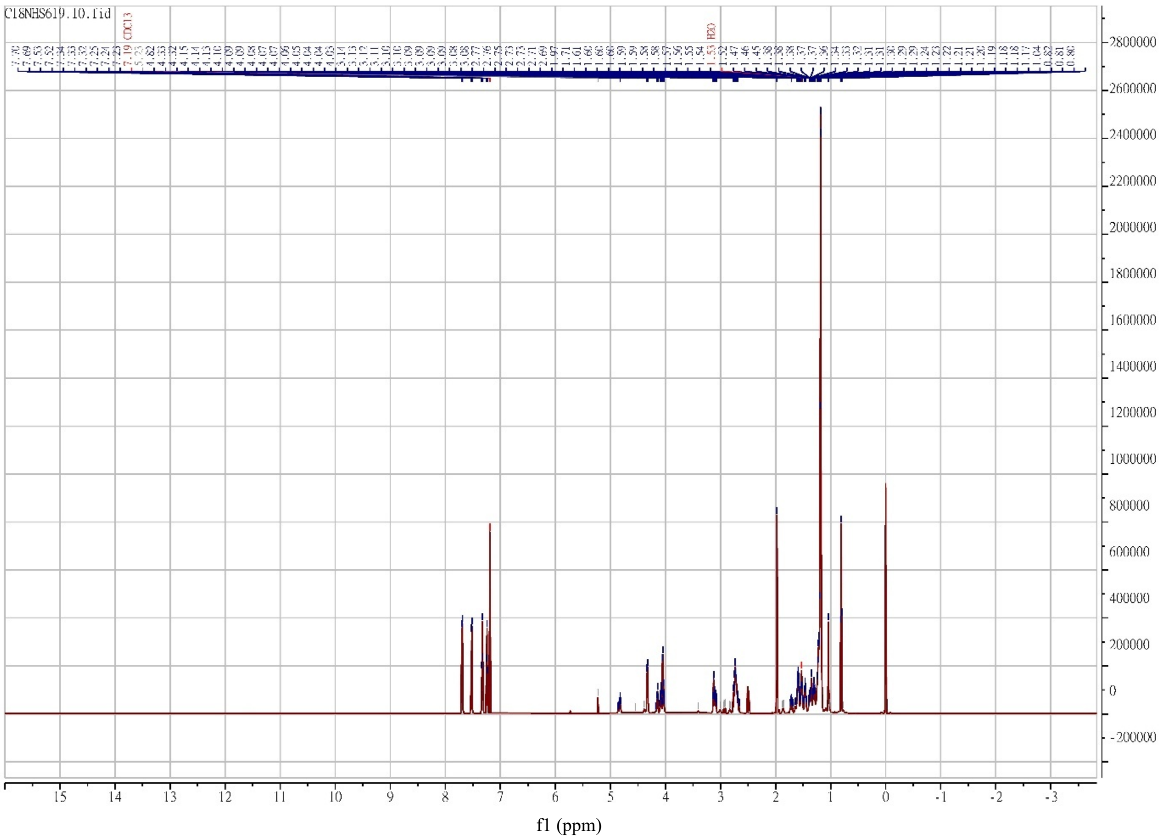

**Figure S9.** $^1H$ NMR spectrum of compound **3**.

#### Compound 3

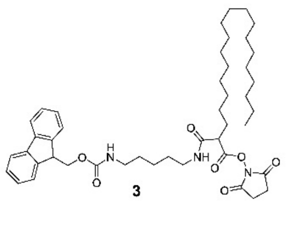


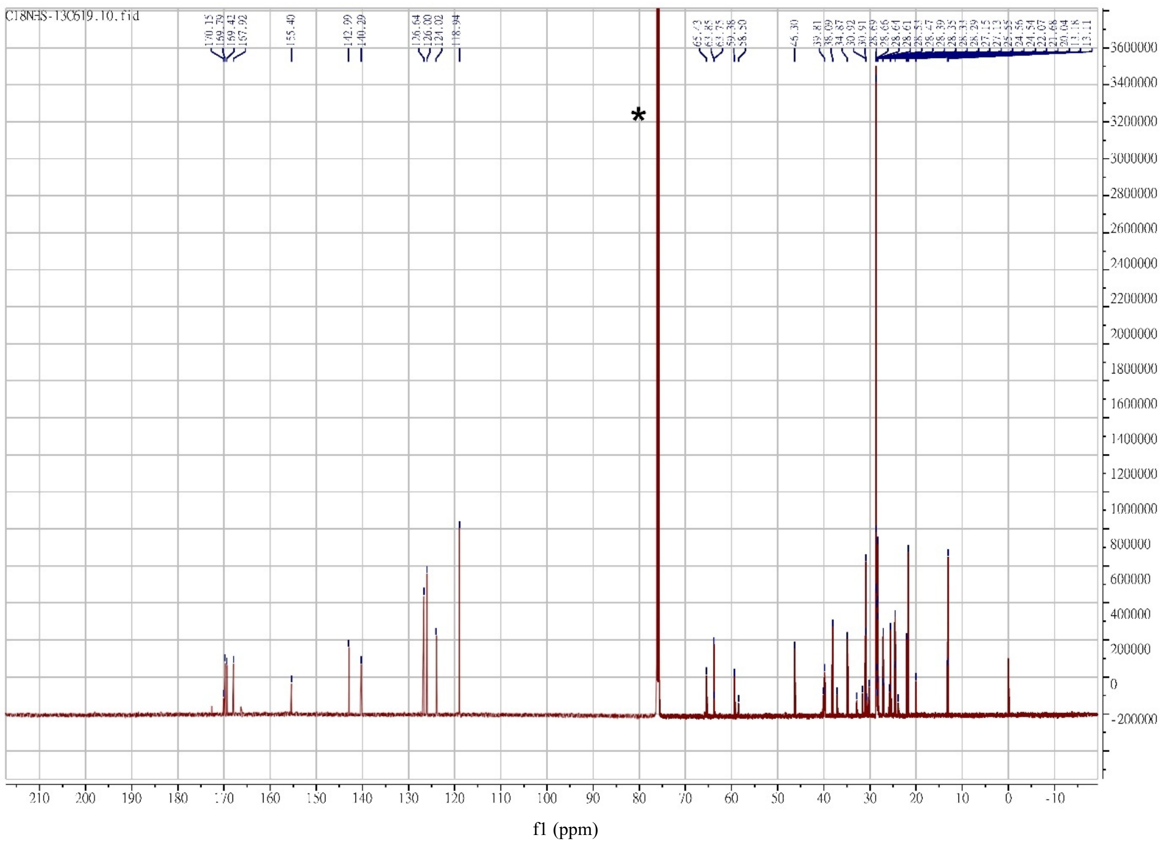

**Figure S10.** $^{13}C$ NMR spectrum of compound **3**.

#### Compound 4

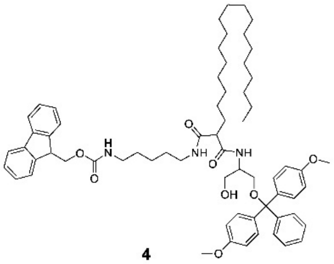

4

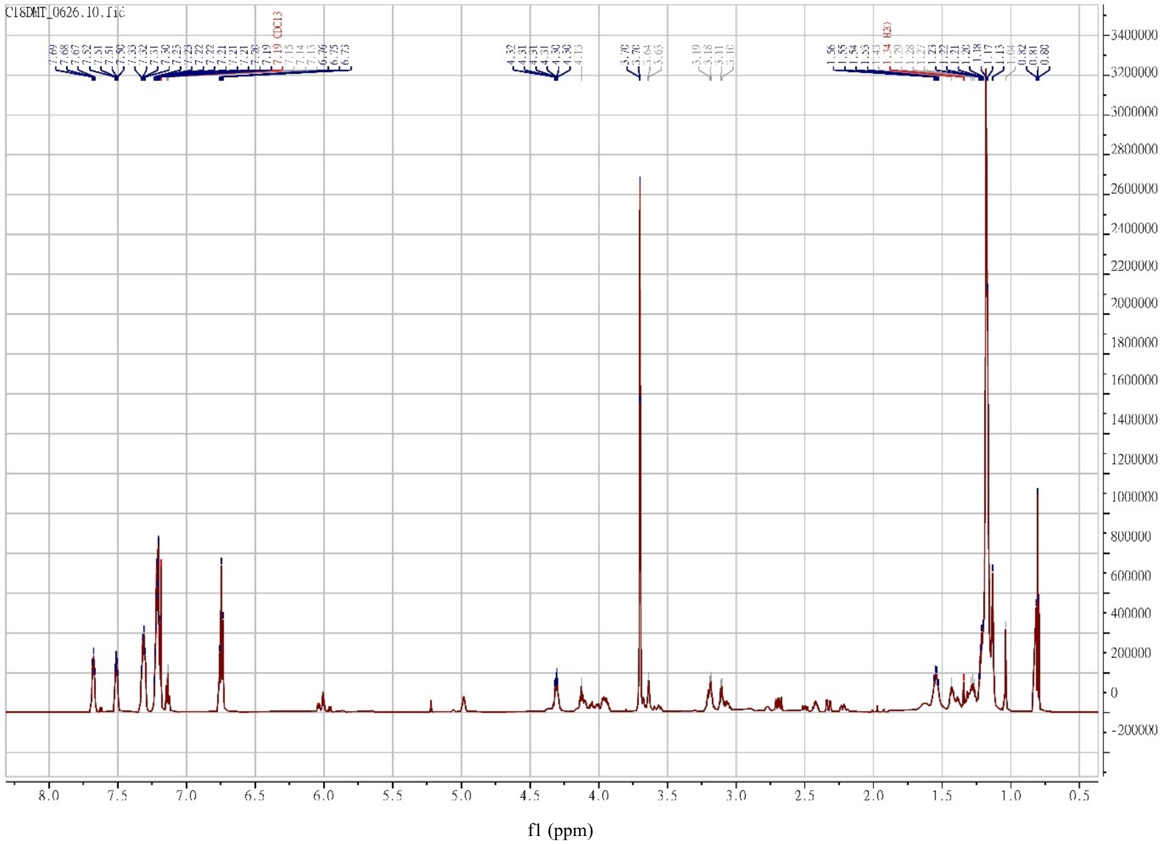

**Figure S11.** $^{1}H$ NMR spectrum of compound **4**.

### Compound 4

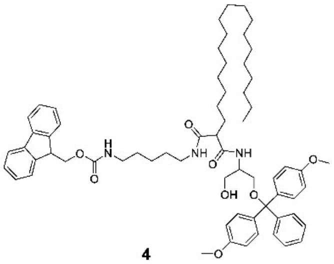

4

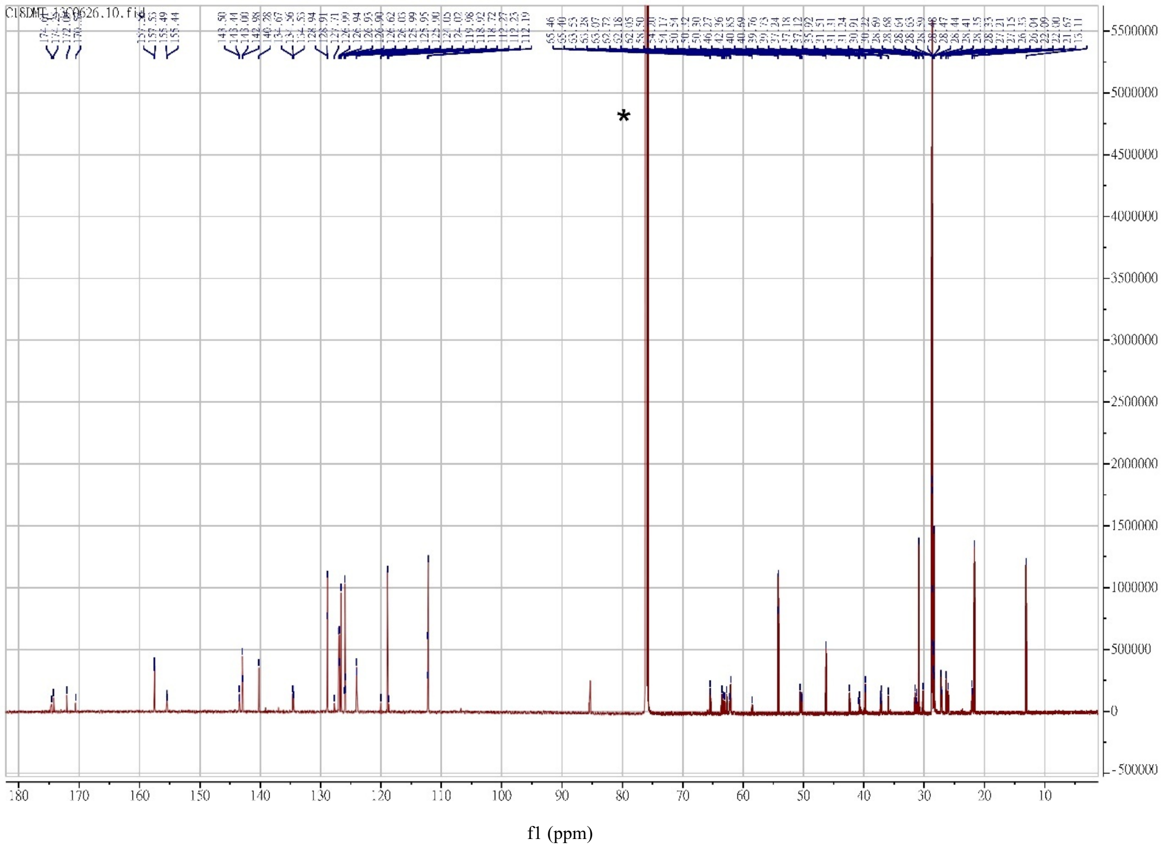

Figure S12. $^{13}$C NMR spectrum of compound **4**.

#### Compound 5

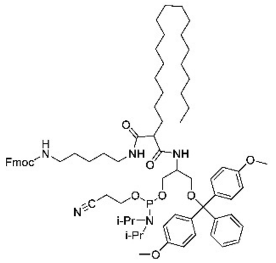


**5**

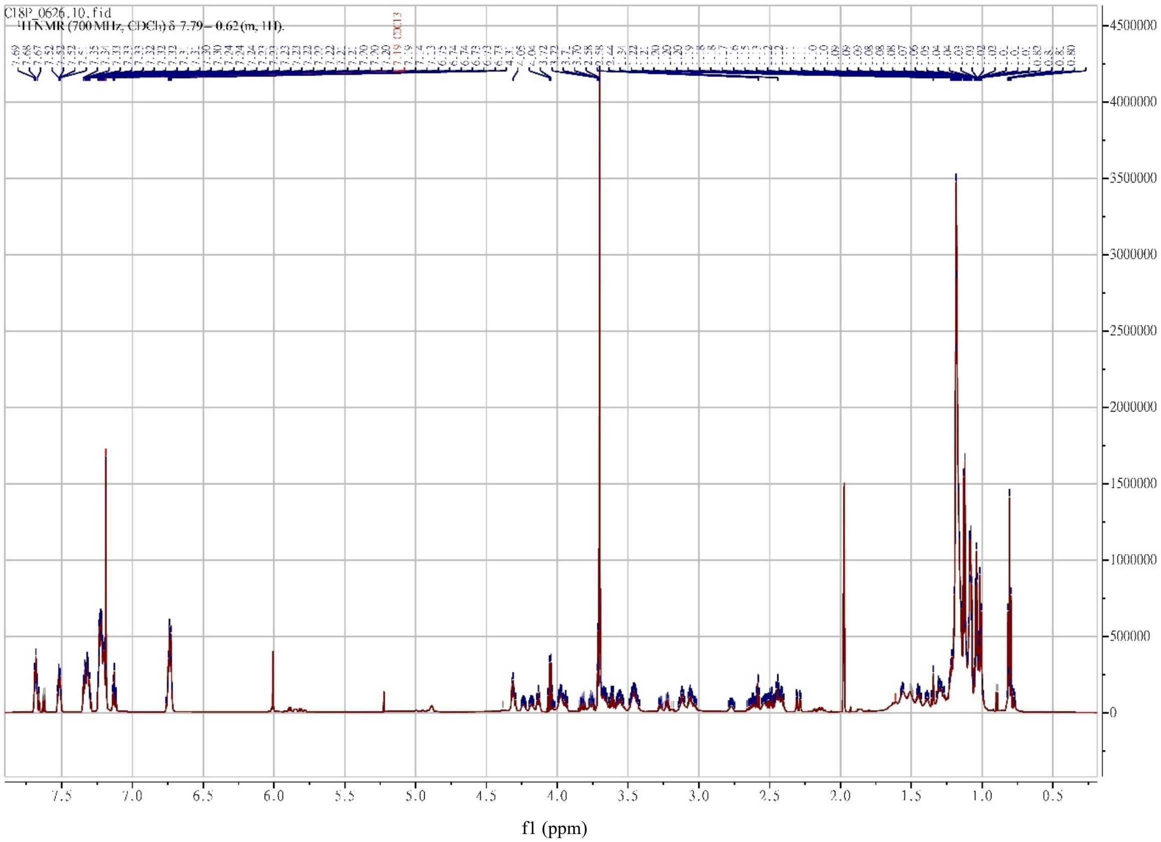


**Figure S13.** $^{1}$H NMR spectrum of compound **5**.

#### Compound 5

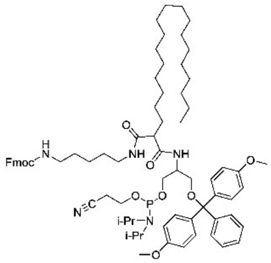


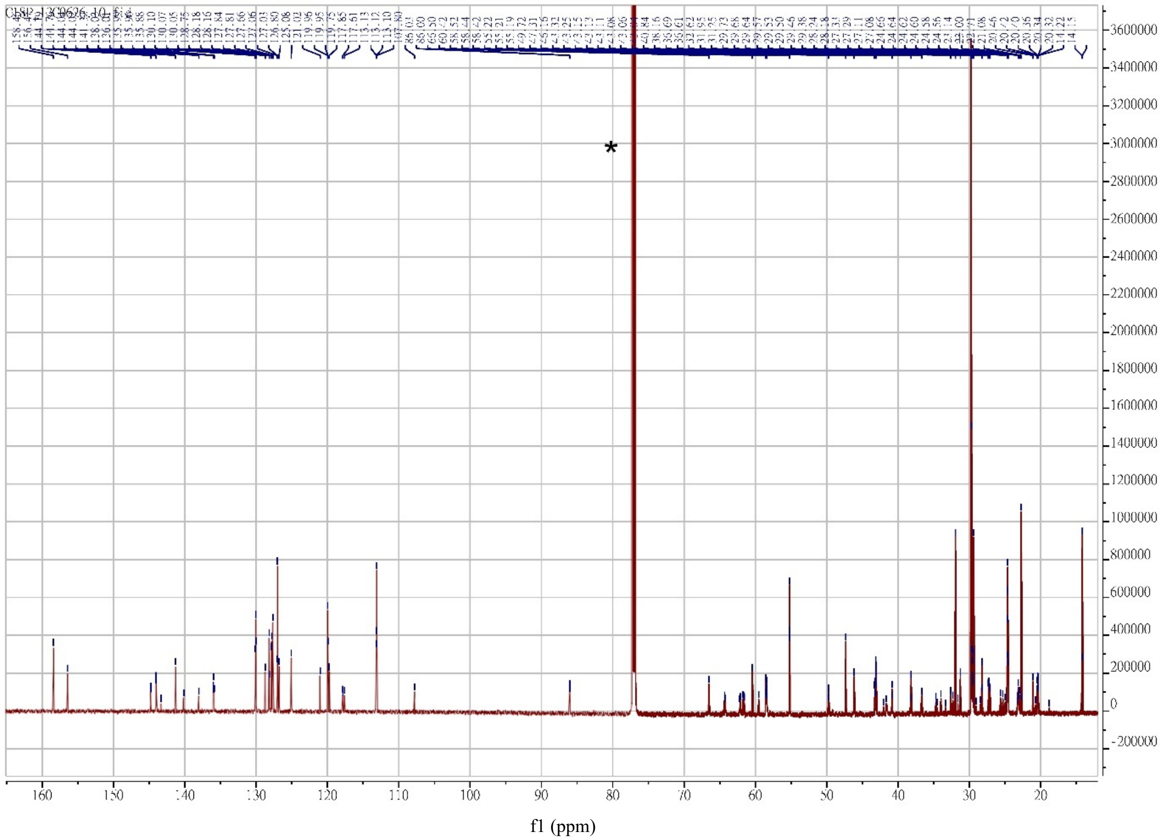


**Figure S14.** $^{13}$CNMR spectrum of compound **5**.

#### Compound 7

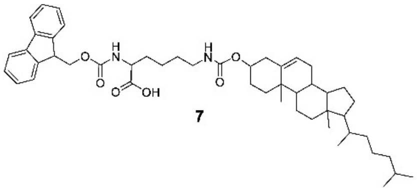

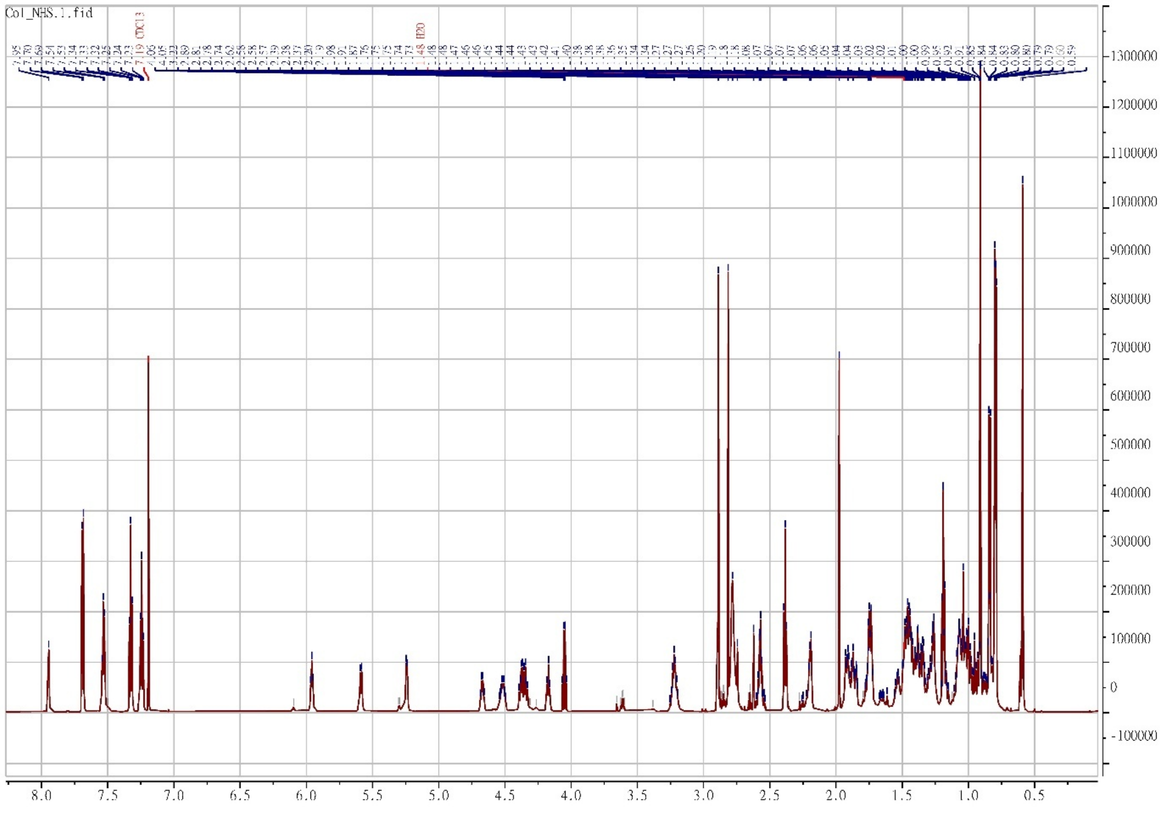


**Figure S15.** $^{1}$H NMR spectrum of compound **7**.

#### Compound 7

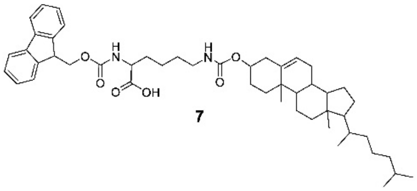
7

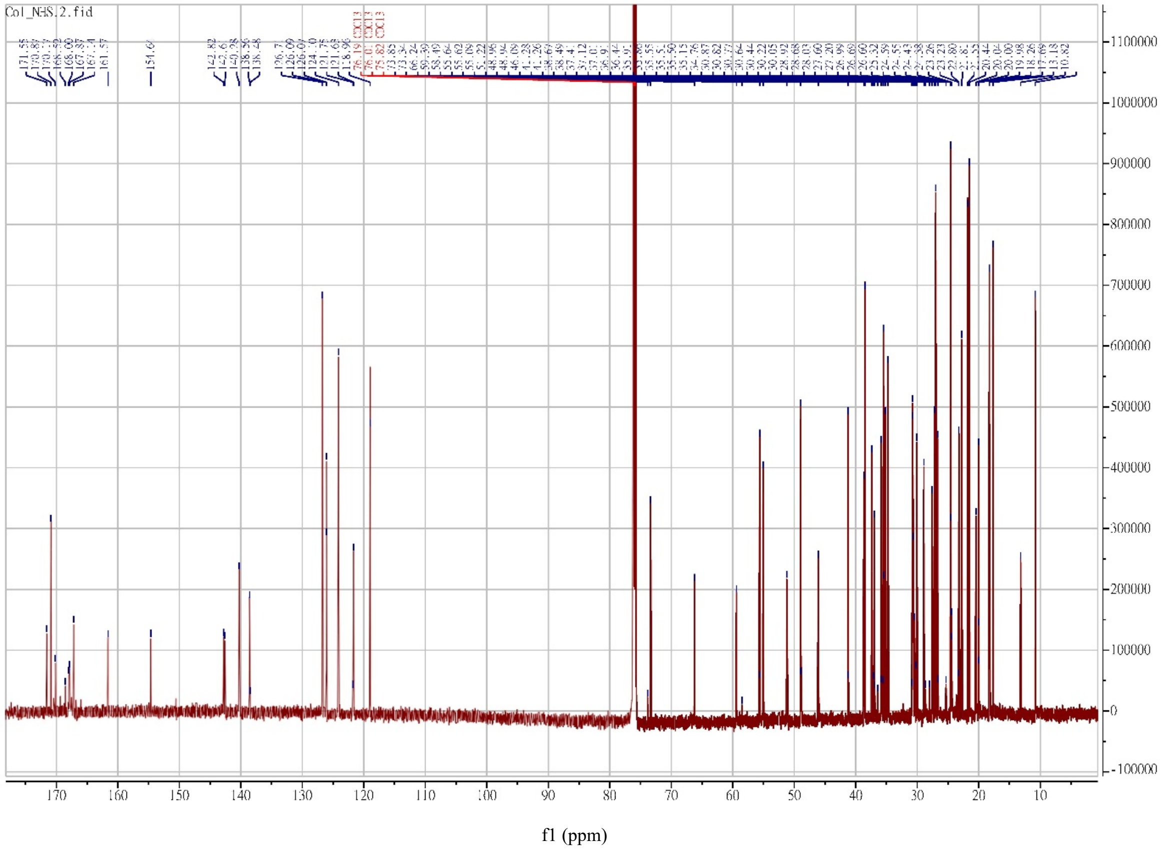

**Figure S16.** $^{13}C$ NMR spectrum of compound **7**.

#### Compound 8

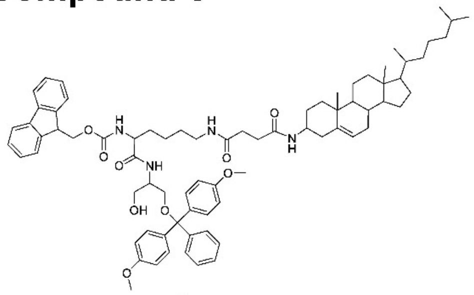


8

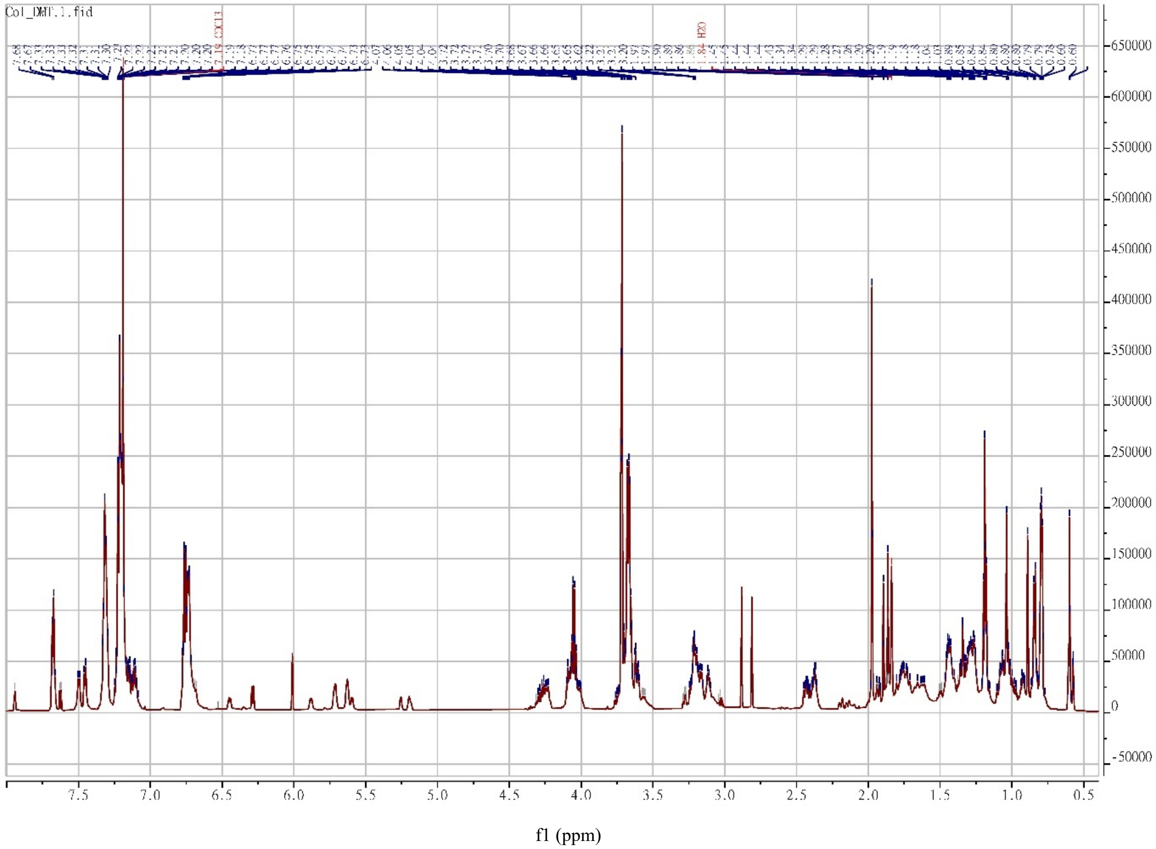


**Figure S17.** $^{1}$H NMR spectrum of compound **8**.

#### Compound 8

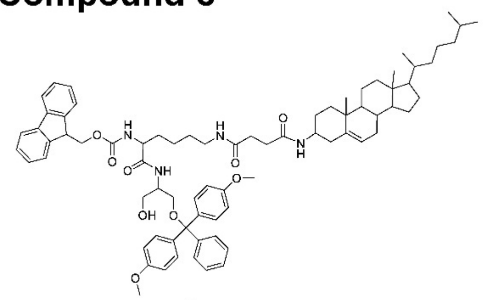

8

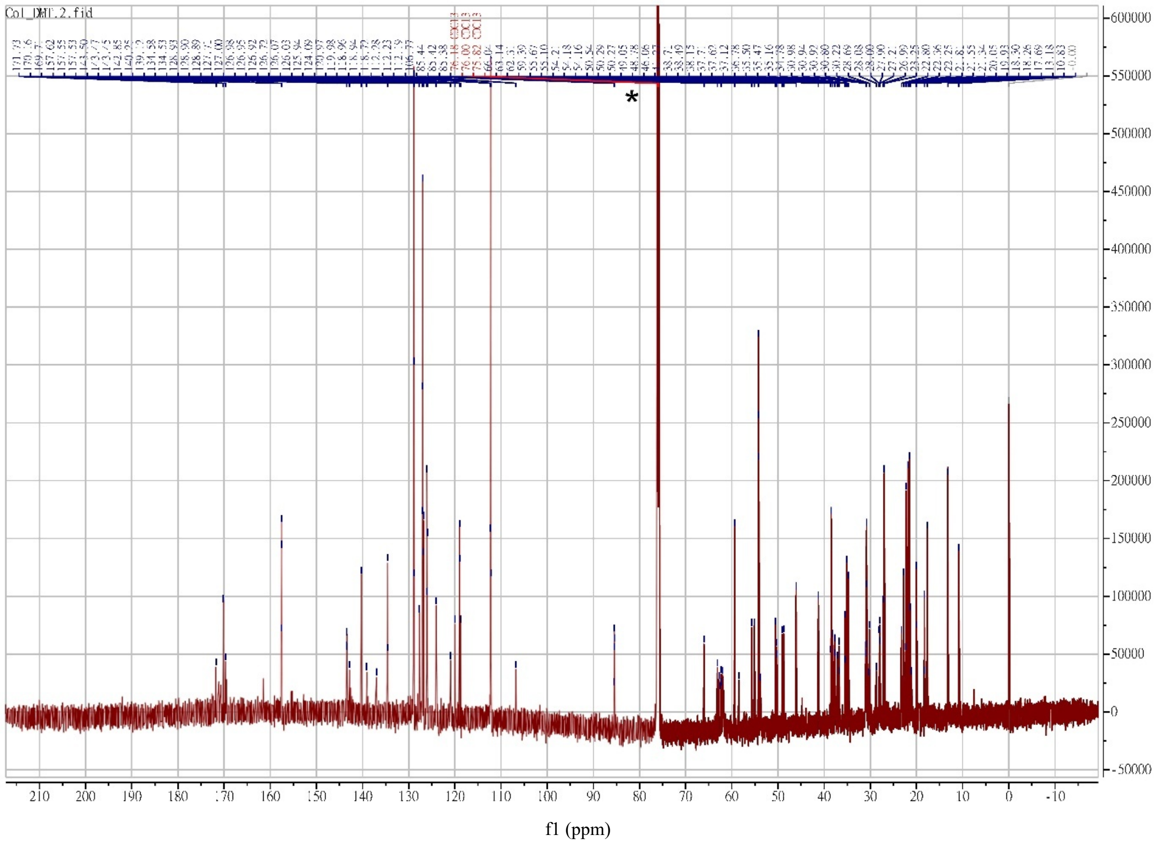


**Figure S18.** $^{13}$C NMR spectrum of compound **8**.

#### Compound 9

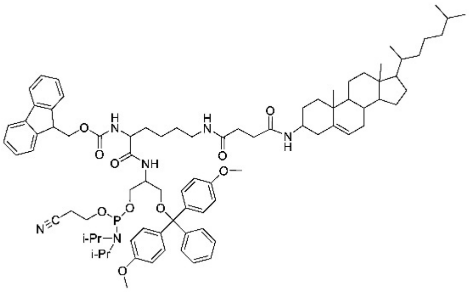

9

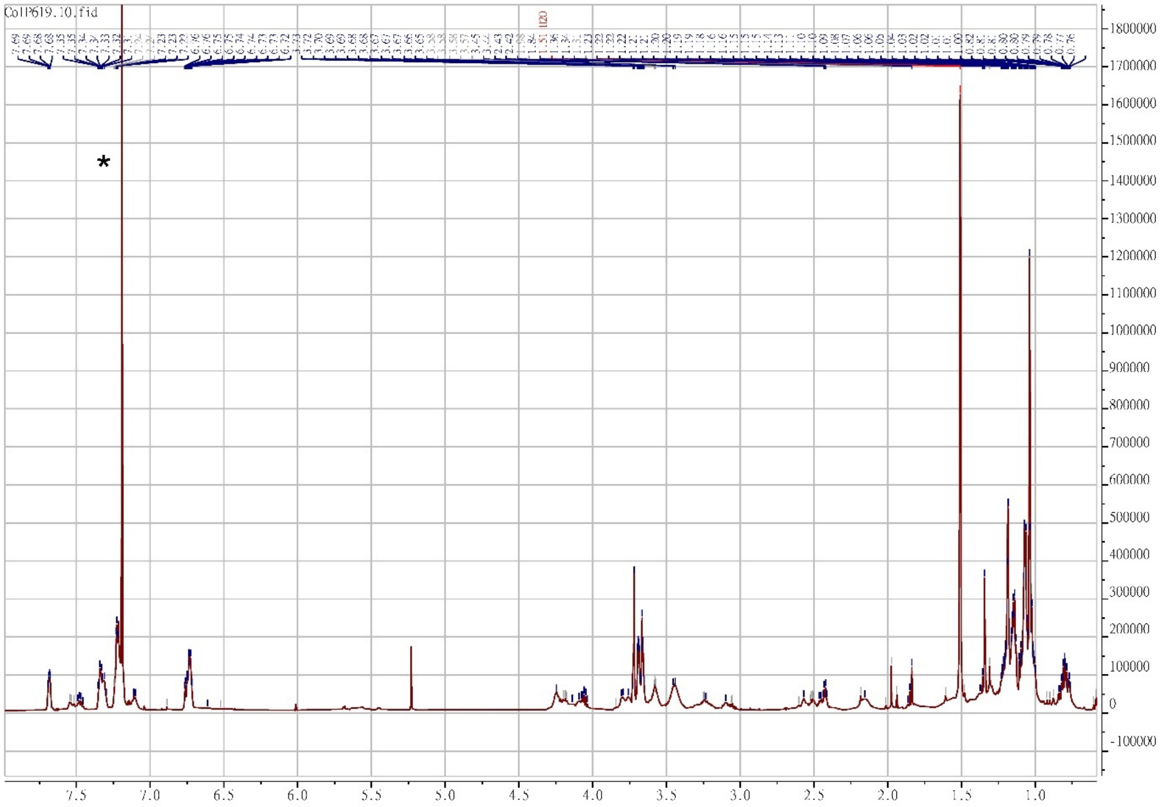


f1 (ppm)

**Figure S19.** $^{1}$H NMR spectrum of compound **9**.

#### Compound 9

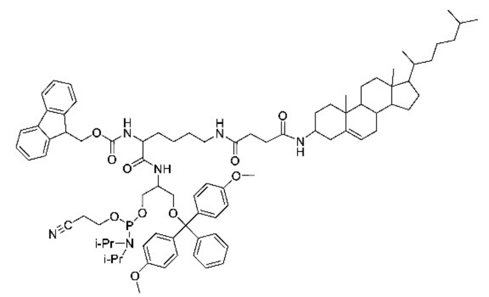

9

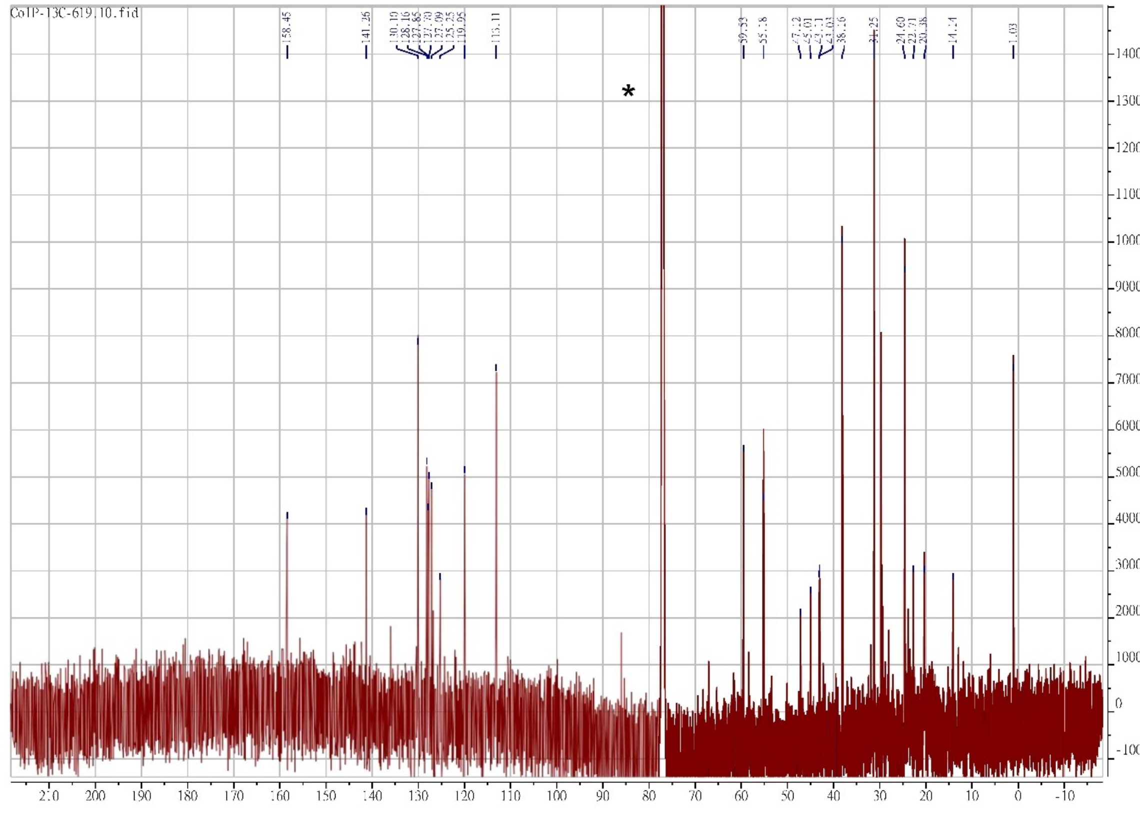


**Figure S20.** $^{13}$C NMR spectrum of compound **9**.

### Compound 10

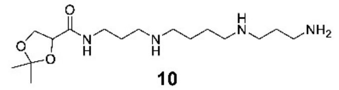


**10**

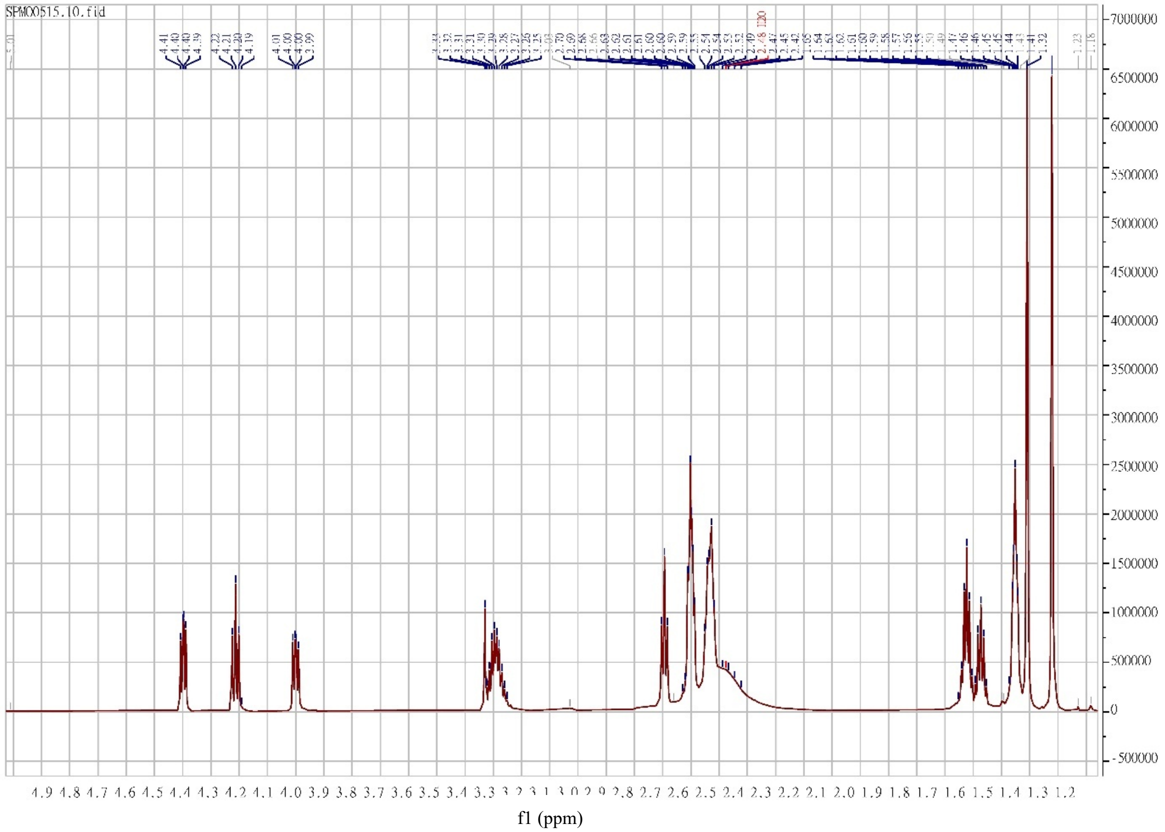

**Figure S21.** [1]H NMR spectrum of compound **10**.

#### Compound 11

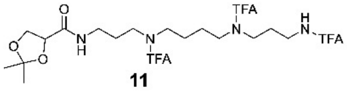


11

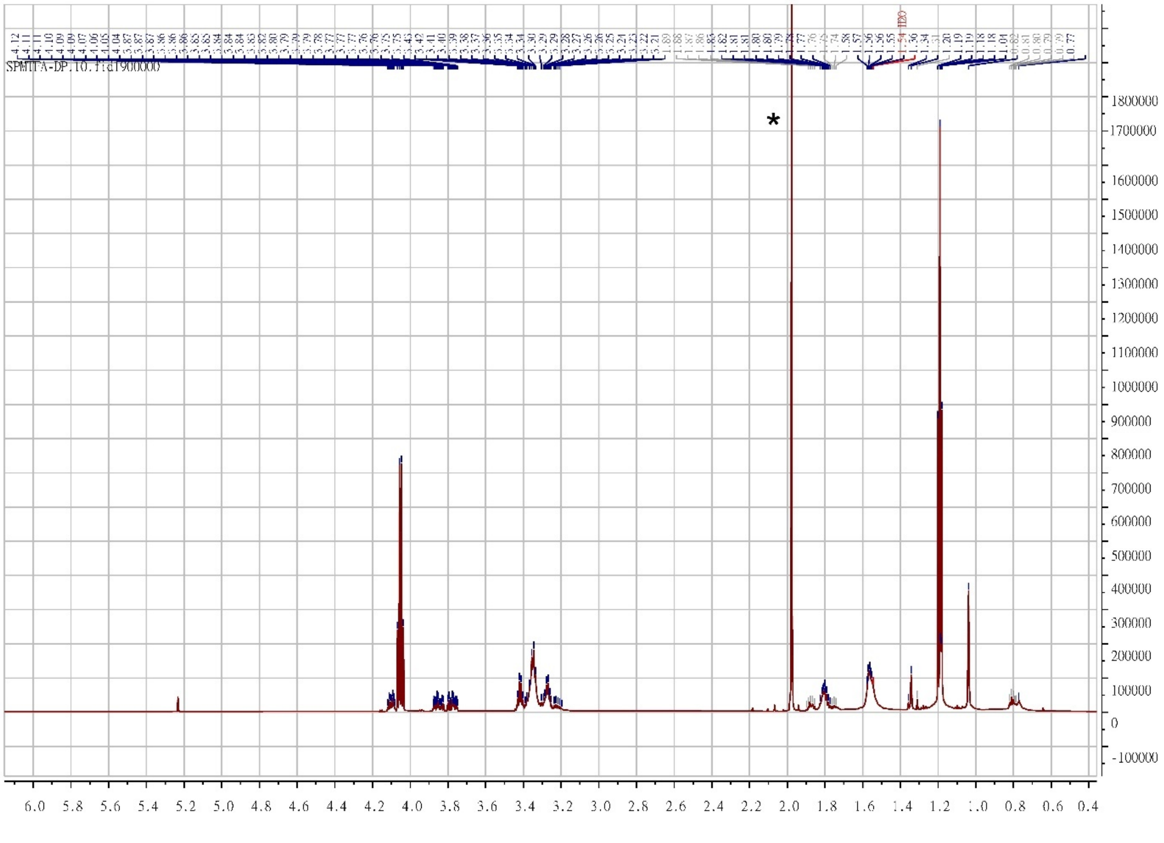


**Figure S22.** $^{1}$H NMR spectrum of compound **11**.

### Compound 11

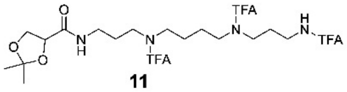

11

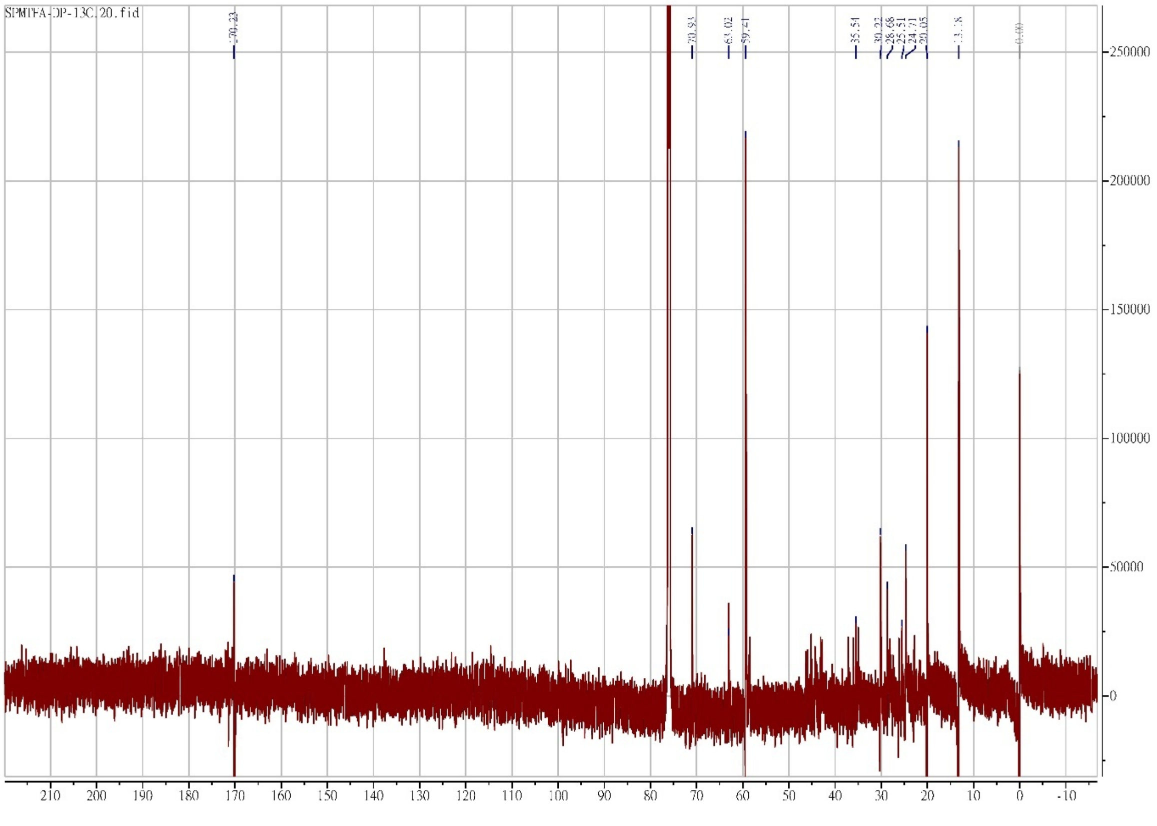


**Figure S23.** $^{13}$C NMR spectrum of compound **11**.

#### Compound 12

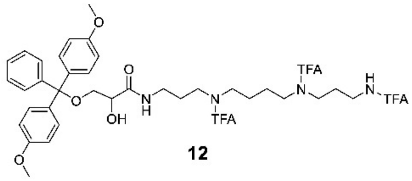


**12**

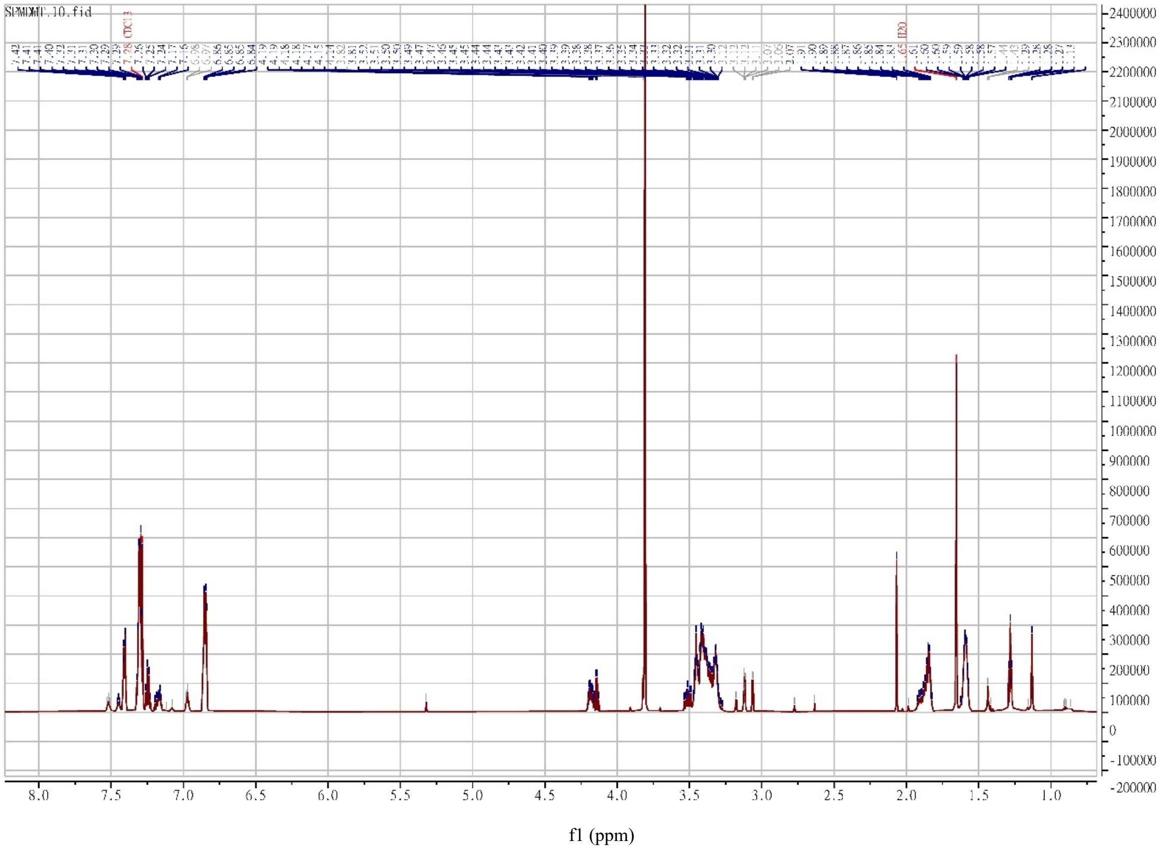


**Figure S24.** $^1$H NMR spectrum of compound **12**.

#### Compound 12

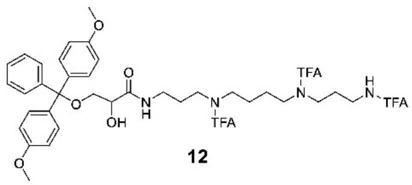


**12**

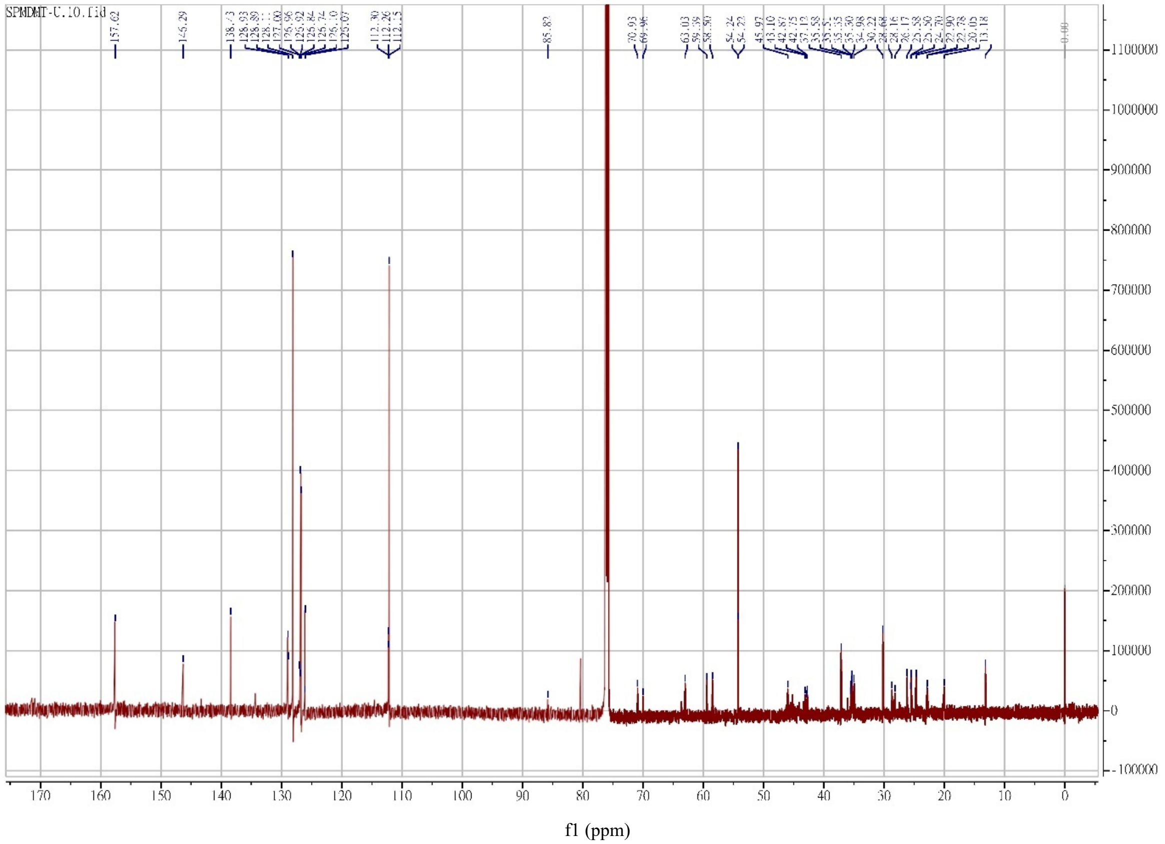


**Figure S25.** $^{13}$C NMR spectrum of compound **12**.

#### Compound 13

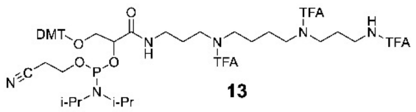


13

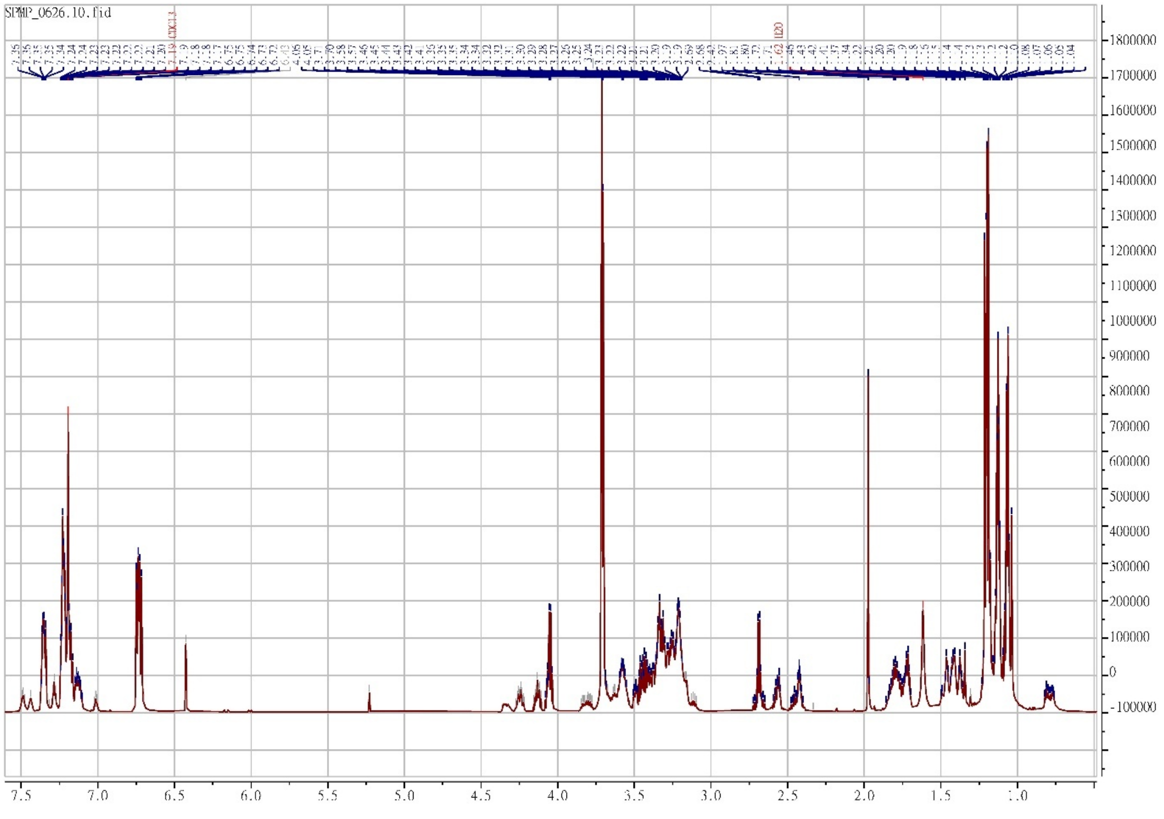


**Figure S26.** $^{1}$H NMR spectrum of compound **13**.

#### Compound 13

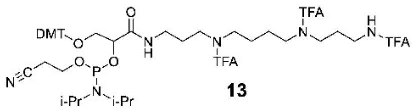

13

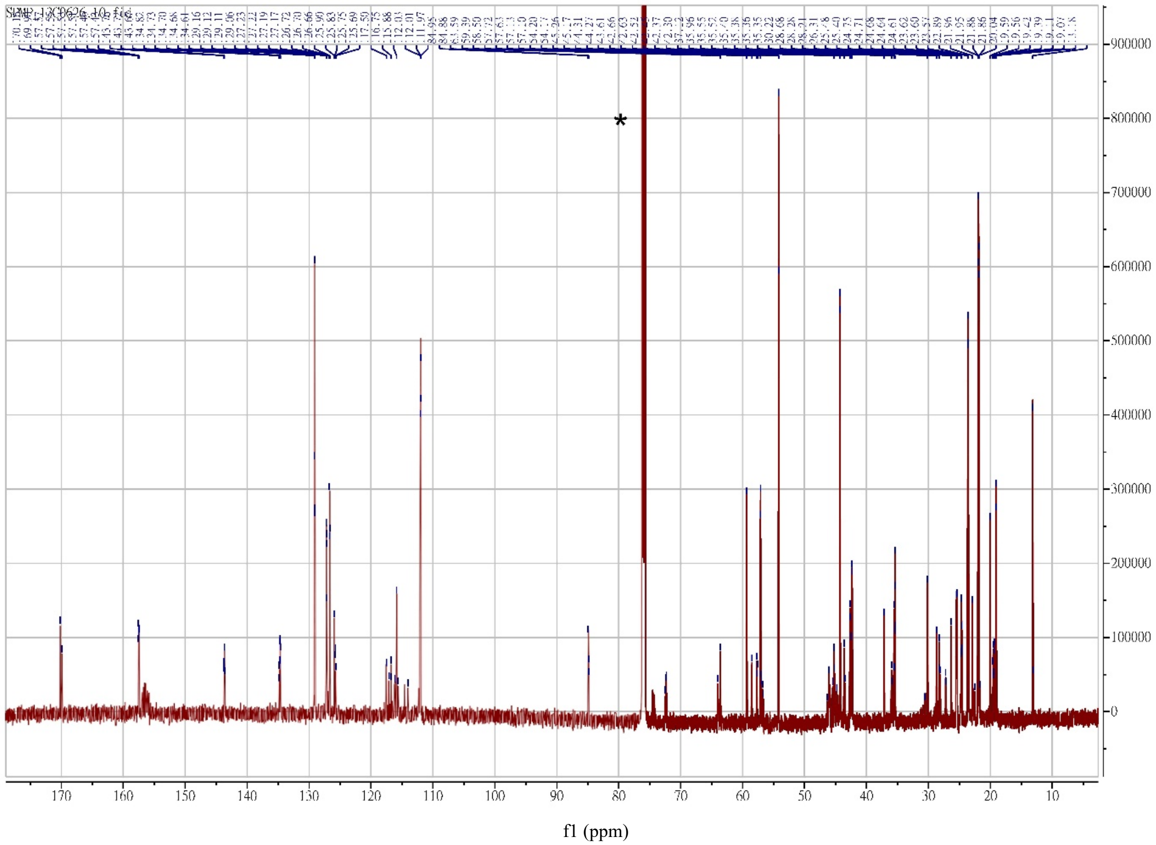


**Figure S27.** $^{13}$C NMR spectrum of compound **13**.

#### Compound 16

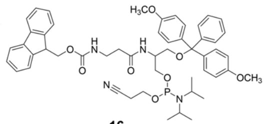


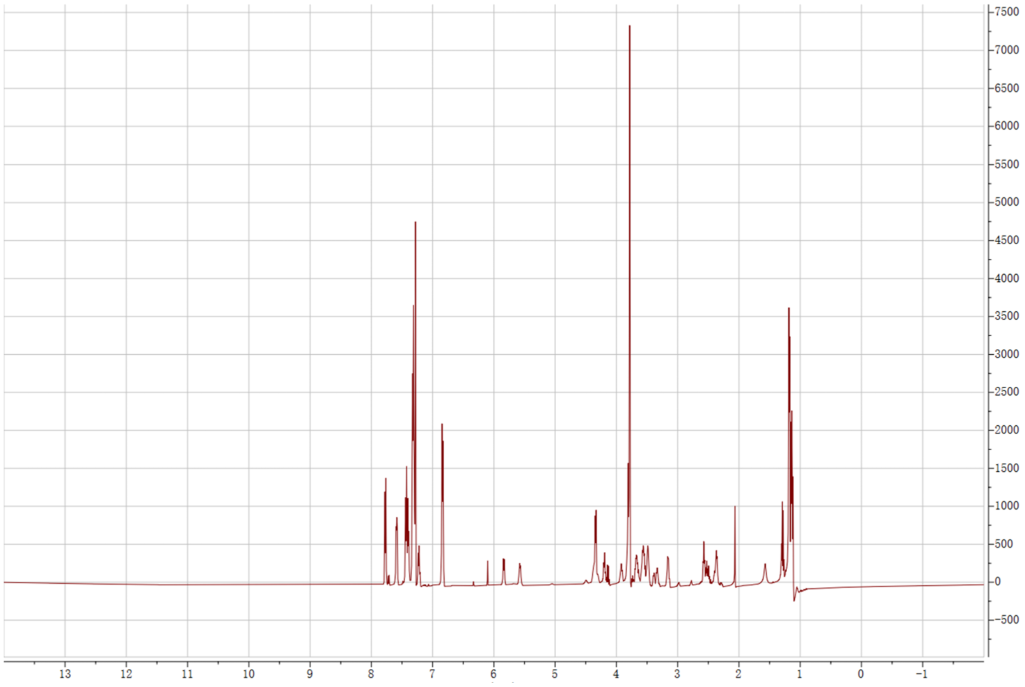


**Figure S28.** [1]H NMR spectrum of compound **16**.

#### Compound 16

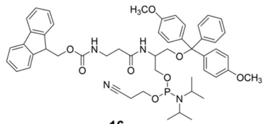

**16**

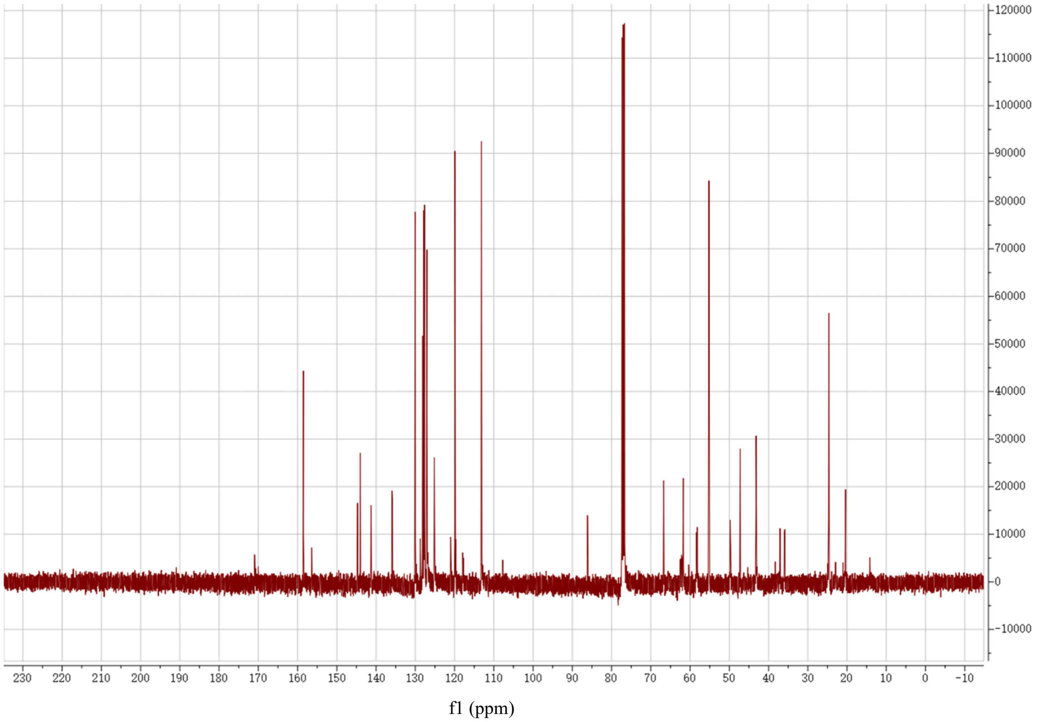


**Figure S29.** $^{13}C$ NMR spectrum of compound **16**.
